# HKU5 bat merbecoviruses use divergent mechanisms to engage bat and mink ACE2 as entry receptors

**DOI:** 10.1101/2025.02.12.637862

**Authors:** Mia Madel Alfajaro, Emma L. Keeler, Ning Li, Nicholas J. Catanzaro, I-Ting Teng, Zhe Zhao, Michael W. Grunst, Boyd Yount, Alexandra Schäfer, Danyi Wang, Arthur S. Kim, Aleksandra Synowiec, Mario A. Peña-Hernández, Ridwan Arinola, Ramandeep Kaur, Bridget L. Menasche, Jin Wei, Gabriel A. Russell, John Huck, Jaewon Song, Aaron Ring, Akiko Iwasaki, Rohit K. Jangra, Sanghyun Lee, David R. Martinez, Walther Mothes, Pradeep D. Uchil, John G. Doench, Alicen B. Spaulding, Ralph S. Baric, Leonid Serebryannyy, Yaroslav Tsybovsky, Tongqing Zhou, Daniel C. Douek, Craig B. Wilen

## Abstract

Identifying receptors for bat coronaviruses is critical for spillover risk assessment, countermeasure development, and pandemic preparedness. While Middle East respiratory syndrome coronavirus (MERS-CoV) uses DPP4 for entry, the receptors of many MERS-related betacoronaviruses remain unknown. The bat merbecovirus HKU5 was previously shown to have an entry restriction in human cells. Using both pseudotyped and full-length virus, we show that HKU5 uses *Pipistrellus abramus* bat ACE2 but not human ACE2 or DPP4 as a receptor. Cryo-electron microscopy (cryo-EM) analysis of the virus-receptor complex and structure-guided mutagenesis reveal a spike and ACE2 interaction that is distinct from other ACE2-using coronaviruses. MERS-CoV vaccine sera poorly neutralize HKU5 informing pan-merbecovirus vaccine design. Notably, HKU5 can also engage American mink and stoat ACE2, revealing mustelids as potential intermediate hosts. These findings highlight the versatility of merbecovirus receptor use and underscore the need for continued surveillance of bat and mustelid species.

## INTRODUCTION

Bats are a major reservoir for coronaviruses with pandemic potential^1,2^. To date, three highly pathogenic betacoronaviruses have spilled over into humans: severe acute respiratory syndrome coronavirus (SARS-CoV), SARS-CoV-2, and Middle East respiratory syndrome coronavirus (MERS-CoV)^3–6^. MERS-CoV, a member of the *Merbecovirus* subgenus, transmits sporadically in humans but has a case-fatality rate of 35%^7^. Hundreds of other bat merbecoviruses circulate in the wild and thus represent a potential pandemic threat^8,9^. However, our ability to risk stratify and develop effective countermeasures against these viruses is precluded by the scarcity of information regarding their receptor use^9^.

Viral receptors mediate host range, tissue tropism, transmission fitness, and pathogenesis^10–12^. Multiple merbecoviruses including MERS-CoV and bat coronaviruses HKU4, HKU25, and 422-CoV use dipeptidyl peptidase-4 (DPP4) as a receptor^13–17^. In contrast, many sarbecoviruses (e.g., SARS-CoV, SARS-CoV-2) as well as the alphacoronavirus HCoV-NL63 use angiotensin converting enzyme 2 (ACE2) for entry^18–21^. The coronavirus spike protein is comprised of S1 and S2 subunits which mediate host receptor attachment and membrane fusion, respectively. S1 contains N-terminal (NTD) and C-terminal domains (CTD) with the CTD containing the classical receptor binding domain (RBD). In addition to receptor binding, cellular proteases such as TMPRSS2 and Cathepsin L prime and activate spike to enable membrane fusion^10,21–23^.

HKU5-like coronaviruses comprise a clade of group 2c merbecoviruses isolated from *Pipistrellus* bats in East Asia^24,25^. Despite being closely related to MERS-CoV and the group 2c bat coronavirus HKU4, the receptor for HKU5 is unknown, hindering initial efforts to rescue a full-length infectious molecular clone of HKU5^9,23,24,26^. Previous studies demonstrated that human cells are permissive to HKU5 but that there was a block to infection at viral entry that could in part be overcome with exogenous trypsin treatment^26,27^. An HKU5 chimera (HKU5-SE) in which the SARS-CoV spike ectodomain replaced that of HKU5 efficiently replicated in human cells and caused severe disease in aged mice^26^. This demonstrates pathogenic potential of the HKU5 viral backbone. African and European bat merbecoviruses (NeoCoV, PDF-2180; MOW15-22, PnNL2018B; respectively) were recently shown to use bat ACE2, albeit by distinct mechanisms^9,28,29^. To date, no merbecoviruses from Asia have been shown to use ACE2 as a receptor. Identifying the receptor for HKU5 and related viruses is crucial for understanding their host range, spillover potential, and pandemic risk.

Here, using a panel of receptor orthologs we identify that a prototypic HKU5 and the related BtPa-BetaCoV/GD2013 (GD2013) use ACE2 from *P. abramus* bats for entry. We characterize this interaction with genetic, biochemical, and structural studies including a 4.2 Å cryo-EM structure of the HKU5^RBD^ and *P. abramus* ACE2 complex. We map key interacting residues via mutagenesis and demonstrate that full-length infectious HKU5 requires *P. abramus* ACE2. We also identify mustelids as possible intermediate hosts for the HKU5 clade. This work will enable improved surveillance and the development of countermeasures for this group of bat coronaviruses.

## RESULTS

### Investigation of receptor usage of diverse bat CoVs

To screen for coronavirus receptors, we synthesized spikes from five diverse bat coronaviruses from three different *Betacoronavirus* subgenera: bat HKU5-LMH03f (*Merbecovirus*, HKU5), *Erinaceus* CoV/2012-74 (*Merbecovirus*, EriCoV/2012), RoBatCoV/HKU9 (*Nobecovirus*, HKU9), RoBatCoV/GCCDC1 (*Nobecovirus*, GCCDC1), and bat Hp-BetaCoV/Zhejiang2013 (*Embecovirus*, HpBeta-CoV) (**Fig. 1a** and Extended Data Fig. 1a). We generated replication defective vesicular stomatitis virus (VSV) pseudovirus expressing each respective spike protein and a Renilla Luciferase (RLuc) reporter. To test whether ACE2 or DPP4 could mediate entry, we initially synthesized expression constructs for 31 DPP4 (**Fig. 1b and Supplementary Table 1**) and 48 ACE2 orthologs (**Fig. 1c and Supplementary Table 2)** from wild, domestic, and peri-domestic species. DPP4 and ACE2 plasmids were transfected into human HEK-293T cells and expression was confirmed by Western blot (Extended Data Fig. 1b,c). First, we investigated whether any DPP4 ortholog(s) could facilitate viral entry of these orphan coronaviruses, in addition to SARS-CoV-2 and MERS-CoV controls. MERS-CoV spike pseudoviruses (VSV-MERS-CoV^spike^-RLuc) readily infected cells expressing human DPP4 as well as other mammalian DPP4 orthologs. However, these DPP4s were not sufficient to promote entry of the other pseudoviruses (**Fig. 1b**). Next, we assessed whether any of these pseudoviruses could efficiently use ACE2 orthologs for entry. The SARS-CoV-2 spike mediated entry of cells expressing the ACE2 of 38/48 ACE2 orthologs, including 19/27 bat species and 19/21 non-bat species, in both HEK-293T (human) and BHK-21 (hamster) cells, the latter of which lacks endogenous ACE2 (**Fig. 1c**). The ability to engage diverse ACE2 orthologs is consistent with the broad host range of SARS-CoV-2^30^. Notably, among the pseudoviruses tested, we detected a significant increase in HKU5 pseudovirus (VSV-HKU5^spike^-Rluc) entry into both *Pipistrellus abramus (P. abramus,* Japanese house bat) and *Neogale vison* (*N. vison,* American mink) ACE2-expressing cells. The four other bat coronavirus spikes did not mediate detectable entry with any tested ACE2 ortholog (**Fig. 1c**). As expected, sequence alignment revealed high sequence variability in the RBDs of ACE2-using and DPP4-using merbecoviruses^31^ (Extended Data Fig. 1d and Extended Data Fig. 2).

**Fig. 1.**
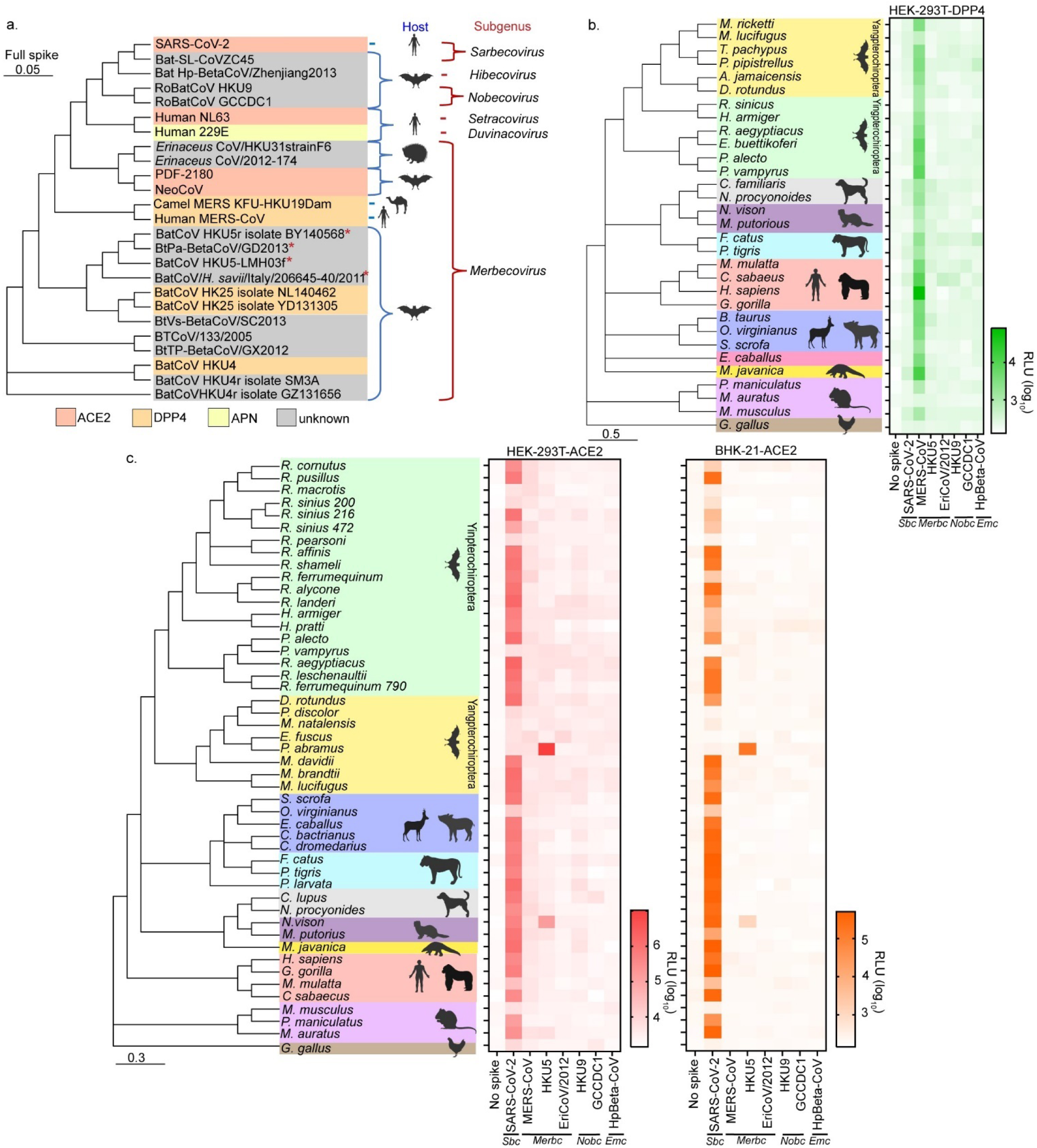
*P. abramus* bat and mink ACE2, but not DPP4, enable HKU5 pseudovirus entry. **a,** Phylogenetic analysis of the full-length amino acid sequences of coronavirus spikes from representative subgenera. Hosts and receptors are indicated. Asterisks (*) highlight bat coronaviruses used in this study. **b,** HEK-293T cells were transiently transfected with the indicated animal DPP4 constructs and infection of each spike pseudovirus was assessed. MERS-CoV pseudovirus uses diverse DPP4 orthologs but other spike pseudoviruses do not use DPP4. **c,** HEK-293T or BHK-21 cells were transiently transfected with the indicated animal ACE2 constructs. Pseudotyped virus infection with coronavirus spikes revealed that HKU5 uses *P. abramus* and *N. vison* ACE2 for entry. In contrast, SARS-CoV-2 uses diverse ACE2 orthologs. The mean of three technical replicates is plotted from one of two independent experiments. Hu, human; RLU, relative light units; CoV, coronavirus; *Sbc*: sarbecovirus; *Merbc*, merbecovirus; *Nobc*, nobecovirus; *Emc*, embecovirus.

### Characterization of bat coronavirus HKU5 utilization of *P. abramus* and *N. vison* ACE2 for entry

Next, we sought to validate *P. abramus* ACE2 and *N. vison* ACE2 as entry receptors for HKU5 from the pseudovirus entry screen. We transfected a targeted panel of ACE2 and DPP4 constructs in HEK-293T cells and used pseudoviruses to assess viral entry. Consistent with our initial screen, HKU5 efficiently used *P. abramus* and *N. vison* ACE2 but not DPP4 for entry (**Fig. 2a,b**). Next, we generated HKU5^spike^ VSV pseudovirus expressing green fluorescence protein (VSV-HKU5^spike^-eGFP) to assess infection on a per cell basis. Consistent with the luciferase results, VSV-HKU5^spike^-eGFP infected HEK-293T cells expressing *P. abramus* ACE2 and *N. vison* ACE2 (**Fig. 2c,d**). At a low multiplicity of infection (MOI), VSV-SARS-CoV-2^spike^-eGFP infected cells expressing human ACE2 and *N. vison* ACE2, but not *P. abramus* ACE2. Next, we tested if murine leukemia virus (MLV) pseudovirus coated with *P. abramus* ACE2 protein (VSV-*P. abramus*^ACE2^-Gluc) could infect HEK-293T target cells expressing HKU5^spike^ on their surface (**Fig. 2e**). MLV bearing *P. abramus* ACE2 readily infected HEK-293T cells in a HKU5^spike^- and ACE2-dependent manner, revealing that HKU5^spike^ and *P. abramus* ACE2 are necessary and sufficient for entry.

**Fig. 2.**
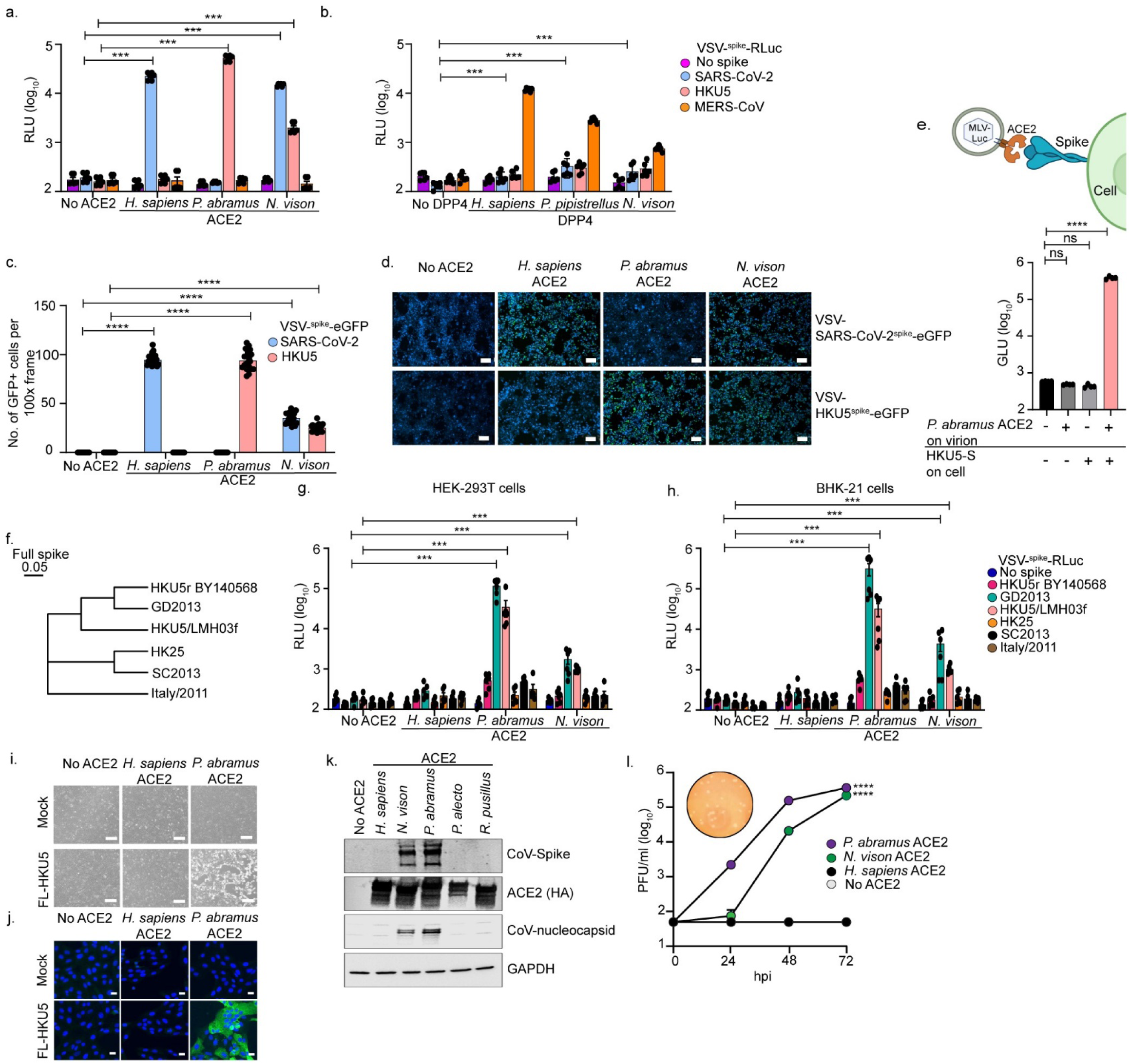
HKU5 uses *P. abramus* bat and mink ACE2 as entry receptors. **a,** HEK-293T cells were transiently transfected with plasmids expressing *H. sapiens* ACE2, *P. abramus* ACE2, *N. vison* ACE2, or a no ACE2 control. Cells were infected with VSV pseudoviruses encoding luciferase and spikes of SARS-CoV-2, HKU5, or MERS-CoV **b,** HEK-293T cells were transiently transfected with plasmids expressing *H. sapiens* DPP4, *P. pipistrellus* DPP4, *N. vison* DPP4, or no DPP4 control. Transfected cells were infected with pseudotyped viruses expressing spike of SARS-CoV-2, HKU5, and MERS-CoV. **c-d,** HEK-293T cells transiently expressing the indicated ACE2s were infected with VSV pseudovirus carrying the enhanced green fluorescent (eGFP) protein. The number of positive eGFP cells was counted in 10 frames at 100x magnification. **e,** Murine Leukemia Virus (MLV) expressing *P. abramus* ACE2 (MLV-*P. abramus*^ACE2^-GLuc) and a Gaussia Luciferase reporter were assessed for the ability to enter HKU5 spike-expressing HEK-293T cells. Data shown is from quadruplicate infections and is representative of two independent experiments. **f,** A phylogenetic tree of full-length spikes related to HKU5. **g,h**, HEK-293T (**g**) and BHK-21 (**h**) cells were transfected with the indicated ACE2s, and pseudotyped virus entry efficiency of HKU5-related viruses was assessed by a Renilla Luciferase assay. **i-l,** Infection in Vero81 cells stably expressing ACE2 constructs and challenged with full-length HKU5 (FL-HKU5). (**i**) Brightfield images from Vero81 cells infected with FL-HKU5 virus. (**j**) FL-HKU5 positive cells were identified by fluorescent microscopy for HKU5 nucleoprotein. HKU5 induces syncytia formation in *P. abramus* ACE2-expressing cells. (**k**) Expression levels of spike, ACE2, nucleocapsid, and GAPDH analyzed by Western blotting at 24 hpi. (**l**) FL-HKU5 can plaque in Vero81/ACE2 cells (inset). Viral kinetics of the FL-HKU5 virus in Vero81/ACE2 cells were analyzed by plaque assay (plaque forming units (PFU)/ml). hu, human; RLU, relative light units. Images in (D) are representative from two independent experiments. Data in (**a, b, g, h**) are pooled from two independent experiments. Scale bars are at 100 µm for panels d and i, and at 50 µm for panel j. Statistical analyses were performed using two-tailed unpaired Student’s *t*-tests and one-way repeated measures ANOVA. Data are represented as mean ± s.e.m. ns, not significant, *\*P<0.05,**P<0.01, ***P<0.001,****P<0.0001*.

To investigate whether other merbecoviruses closely related to the prototypic HKU5 isolate LMH03f could use *P. abramus* and *N. vison* ACE2, we synthesized five additional bat coronavirus spike proteins: BatCoV HKU5r isolate BY140568 (HKU5r BY140568), GD2013, BatCoV HKU25 isolate NL140462 (HKU25), BtVs-BetaCoV/SC2013 (SC2013), and BatCoV/H.savii/Italy/206645-40/2011 (Italy/2011)^32–36^ (**Fig. 2f**). HKU5r BY140568, GD2013, and HKU25 were isolated from *Pipistrellus* bats while SC2013 and Italy/2011 were isolated from *Vespertilio superans* and *Hypsugo savii* bats, respectively^32–36^. GD2013 could use both *P. abramus* and *N. vison* ACE2 for entry while the four other coronaviruses could not use any of the tested ACE2 orthologs (**Fig. 2g,h**). Together this confirms that HKU5 and GD2013 use *P. abramus* and *N. vison* ACE2 as receptors, while the receptors for genetically similar coronaviruses remain unknown.

To confirm the pseudovirus data with full-length infectious HKU5, we inoculated Vero-CCL-81 (Vero81) cells stably expressing *P. abramus* ACE2 (Vero81/*P. abramus* ACE2) with a full-length HKU5 infectious recombinant virus (FL-HKU5). We observed virus-induced cytopathic effects at 72 hours post-infection (hpi) in Vero81 cells expressing *P. abramus* ACE2 but not human ACE2 (**Fig. 2i**). FL-HKU5 readily infected Vero81/*P. abramus* ACE2 cells as demonstrated by the presence of nucleoprotein antigen-positive cells and syncytia formation at 24 hpi (**Fig. 2j**). Next, we challenged HEK-293T cells transiently expressing ACE2 orthologs with FL-HKU5. The HKU5 spike and nucleocapsid proteins were detected by Western blot from cells expressing *P. abramus* ACE2 and *N. vison* ACE2 but not in cells expressing human or other bat (*Pteropus alecto* and *Rhizomucor pusillus*) ACE2s (**Fig. 2k**). Next, we performed a viral growth curve with FL-HKU5 on Vero81 cells stably expressing human ACE2, *P. abramus* ACE2, or *N. vison* ACE2. Vero81/*P. abramus* ACE2 cells infected with FL-HKU5 exhibited a greater than one-log increase in virus titer at 24 hpi and nearly four-log increase in viral titers by 72 hpi (**Fig. 2l**). In contrast, the viral titers in Vero81/*N. vison* ACE2 cells did not show an increase at 24 hpi, but rapidly caught up by 48 hours, resulting in a nearly four-log increase by 72 hpi (**Fig. 2l**). These findings indicate that *P. abramus* ACE2 or *N. vison* ACE2 expression is sufficient to facilitate productive infection and spread of native HKU5 (**Fig. 2l**).

### Evaluation of HKU5 receptor binding domain for targeted ACE2 binding

Next, we sought to determine the mechanism by which HKU5^spike^ interacts with ACE2. We generated mouse immunoglobin G (IgG) Fc-fusion proteins containing the RBD (RBD-Fc) from either SARS-CoV2 (SARS-CoV-2^RBD^-Fc), MERS-CoV (MERS-CoV^RBD^-Fc), PDF-2180 (PDF-2180^RBD^-Fc), or HKU5 (HKU5^RBD^-Fc) spike proteins (Extended Data Fig. 3a,b). We transiently expressed human ACE2, *P. abramus* ACE2, *N. vison* ACE2, or human DPP4 on HEK-293T cells and assessed RBD-Fc binding to these cells by flow cytometry. SARS-CoV-2^RBD^-Fc readily bound human and *N. vison* ACE2 but not *P. abramus* ACE2 or human DPP4 (**Fig. 3a**), consistent with our data and prior studies^35^. MERS-CoV^RBD^-Fc bound human DPP4 but not to any tested ACE2 ortholog (**Fig. 3b**). The PDF-2180^RBD^-Fc bound to *P. abramus* and *N. vison* ACE2 but exhibited low levels of binding to human ACE2 (**Fig. 3c**). The HKU5^RBD^-Fc showed robust binding to cells displaying *P. abramus* ACE2 (**Fig. 3d**). However, binding of HKU5^RBD^-Fc to *N. vison* ACE2 was below the limit of detection (**Fig. 3d**).

**Fig. 3.**
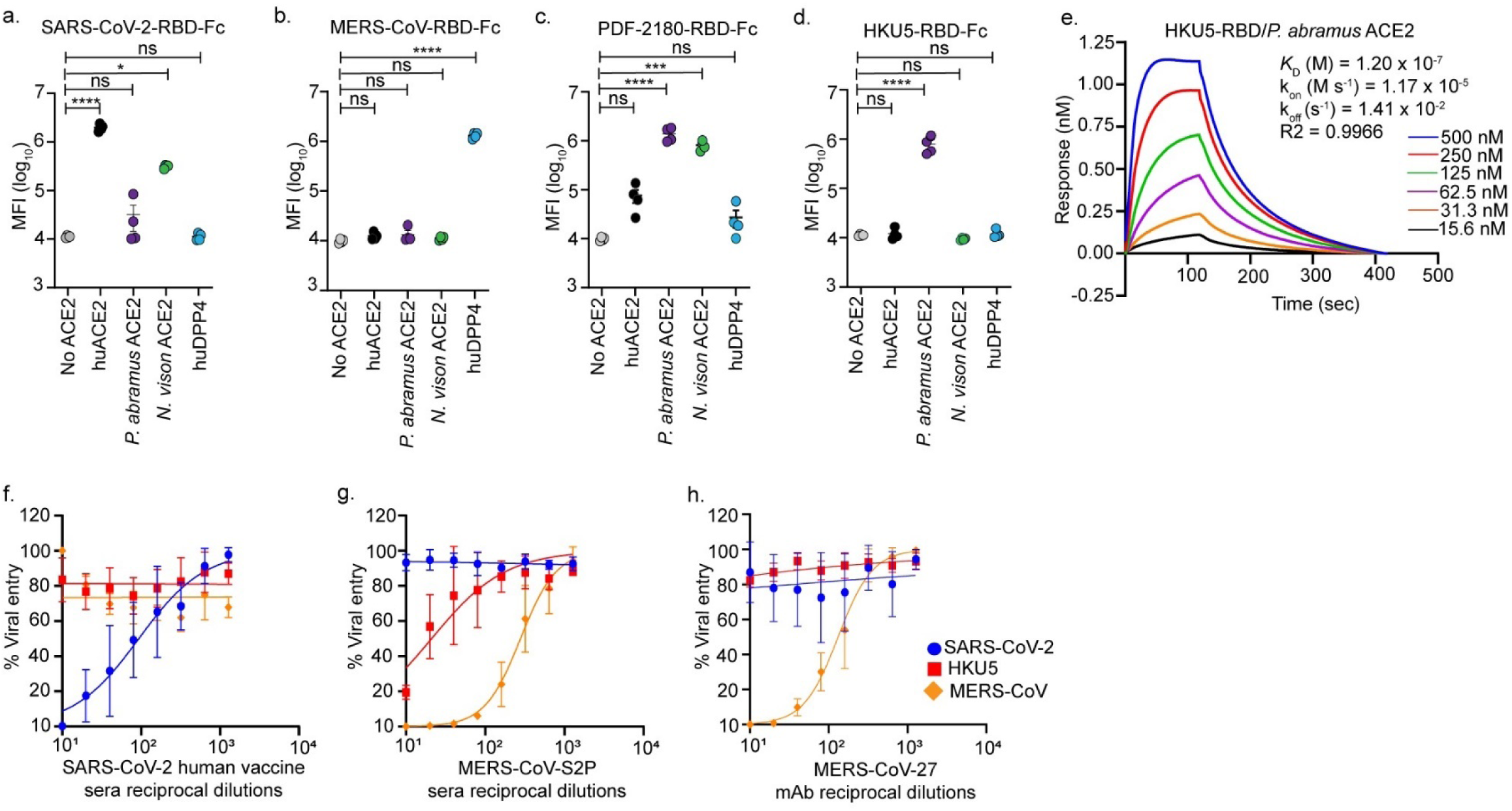
The receptor binding domain of HKU5 spike protein is required for species specific ACE2 binding. **a-d**, Flow cytometry analysis of SARS-CoV-2^RBD^-Fc (**a**), MERS-CoV^RBD^-Fc (**b**), PDF-2180^RBD^-Fc (**c**), and HKU5^RBD^-Fc fusion protein (**d**) binding to HEK-293T cells expressing the indicated receptors. The mean fluorescence intensity (MFI) was calculated. HKU5^RBD^-Fc directly binds the *P. abramus* ACE2. **e,** BLI analysis reveals binding kinetics of HKU5^RBD^ with *P. abramus* ACE2. The reported KD values correspond to avidities due to the use of dimeric ACE2 constructs **f-h,** Neutralization assays using SARS-CoV-2 vaccine sera, MERS-CoV-S2P mouse vaccine sera, and a monoclonal antibody against MERS-CoV (MERS-CoV-27) in BHK-21 cells revealed that HKU5 is resistant to SARS-CoV-2 and MERS-CoV elicited antibodies. Statistical analyses were performed using two-tailed unpaired Student’s *t*-tests and one-way ANOVA. Data are represented as mean ± s.e.m. ns, not significant, *\*P<0.05,**P<0.01, ***P<0.001,****P<0.0001*.

To quantify the affinity and kinetics of the spike-receptor interactions, we performed biolayer interferometry (BLI). Recombinant HKU5^RBD^ bound to recombinant *P. abramus* ACE2, with a binding affinity (Kd) of 120 nM (**Fig. 3e**), whereas binding to *N. vison* ACE2 was below the limit of detection (Extended Data Fig. 3c). Consistent with the flow cytometry binding assay, SARS-CoV-2^RBD^ bound to *N. vison* ACE2 (Kd = 1.5 µM) while SARS-CoV-2^RBD^ binding to *P. abramus* ACE2 was undetectable (Extended Data Fig.3 **d,e**). These results demonstrate direct physical interaction between the RBD of HKU5^spike^ and *P. abramus* ACE2.

The identification of *P. abramus* ACE2 as a receptor for HKU5 enables assessment of antibody-mediated protection. To evaluate antibody mediated cross-protection between HKU5, MERS-CoV, and SARS-CoV-2, we tested neutralizing activity of SARS-CoV-2 human vaccine sera, MERS-CoV-S2P mouse vaccine sera, and a MERS-CoV-targeting monoclonal antibody (MERS-27 mAb)^37,38^ against VSV-HKU5^spike^-RLuc (**Fig. 3f-h**). SARS-CoV-2 vaccine sera nor the MERS-27 mAb neutralized VSV-HKU5^spike^-RLuc (**Fig. 3f-h**). The MERS-CoV-S2P mouse sera had detectable but limited neutralizing activity against VSV-HKU5^spike^-RLuc (**Fig. 3g**). Our results highlight antigenic differences between HKU5, SARS-CoV-2, and MERS-CoV and underscore the need for novel mAbs and vaccines that cross-react with HKU5 viruses.

### Structural basis of HKU5 recognition of *P. abramus* ACE2

To understand HKU5^RBD^ engagement with *P. abramus* ACE2, we performed cryo-EM analysis of HKU5^RBD^ bound to dimeric *P. abramus* ACE2. Data processing yielded a structure at 4.2 Å resolution revealing the ACE2 dimer with HKU5^RBD^ bound to each peptidase domain (**Fig. 4a**, Extended Data Fig. 4, Extended Data Fig. 5, **and Supplementary Table 3**). The binding interface buried a surface of ∼1000 Å^2^ on both sides of the interaction area (**Fig. 4b and Supplementary Table 4**). The 1040 Å^2^ ACE2 binding surface on HKU5^RBD^ is formed by regions on α-helices 4 and 5, β strands 5, 6 and 7, and the loop region exiting β5 (**Fig. 4b**). HKU5^RBD^ binds to ACE2 on a surface boarded by the inner side of helix α1, tip of helix α3, glycan on N387 and the R328 helix (**Fig. 4b**).

**Fig. 4.**
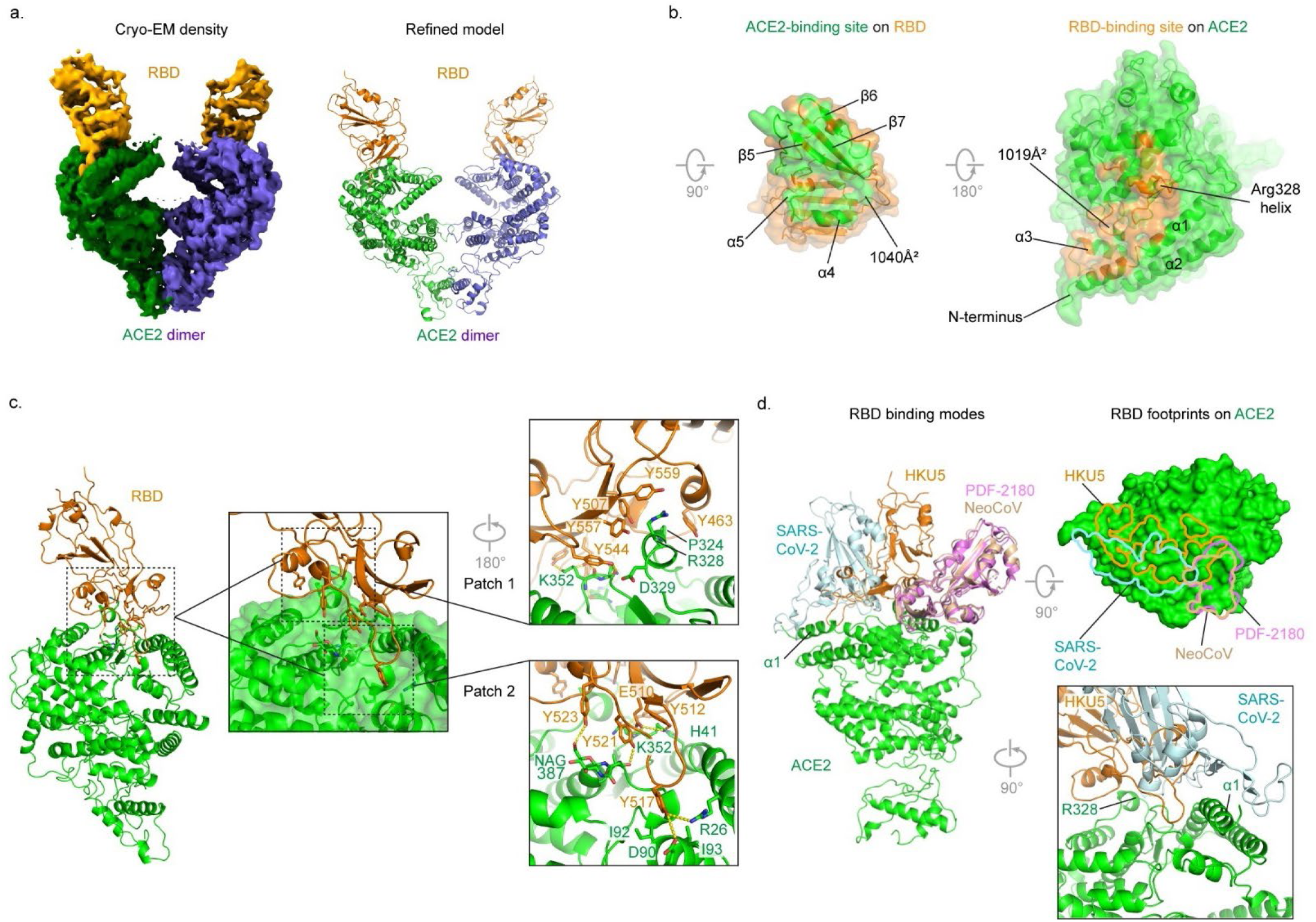
Cryo-EM structure of the HKU5 RBD in complex with *P. abramus* ACE2. **a**, Cryo-EM density (left) and refined model (right) for the HKU5^RBD^ in complex with *P. abramus* ACE2. The ACE2 is colored in green and the RBD is colored in orange. **b,** Footprints of the *P. abramus* ACE2 and HKU5^RBD^ complex are shown in open-book view. **c,** Interactions at the interface. The complex is shown in the 45-degree rotated view compared to the orientation in (a). The interactions are divided in two patches. HKU5^RBD^ tyrosines provide more than 500 Å^2^ binding surface for the *P. abramus* ACE2. Tyrosines in patch 1 form extensive interaction to a helical region on ACE2 that spanning P324^ACE2^, R328^ACE2^ and D329^ACE2^. Patch 2 interactions involve hydrogen bonding between HKU5^RBD^ tyrosines and A385^ACE2^, glycan 387^ACE2^, K352^ACE2^ as well salt bridge between E510^RBD^ and K352^ACE2^. The Y517^RBD^ is wedged between I92^ACE2^ and I93^ACE2^ and interact with R26^ACE2^ and D90^ACE2^. **d,** Comparison of binding modes of HKU5, SARS-CoV-2, NeoCoV and PDF-2180 CoV-RBDs to ACE2. All structures (PDB ID: 6M0J, 7WPO and 7WPZ) were superimposed on to the HKU5^RBD^-bound *P. abramus* ACE2. Footprints of the HKU5^RBD^, SARS-CoV-2^RBD^, PDF2180^RBD^ and NeoCoV^RBD^ are marked on the surface representation of *P. abramus* ACE2 and colored in the same color of respective RBDs in the left panel. A zoom-in view is shown to the lower right to compare the binding difference of HKU5-RBD and SARS-CoV-2^RBD^ relative to the ACE2 α1 helix.

HKU5^RBD^ and *P. abramus* ACE2 form potential hydrogen bonds, salt bridges, polar and stacking interactions (**Supplementary Table 4**). These interactions can be categorized into two major patches. Patch 1 involves five tyrosine residues (Y463^RBD^, Y507^RBD^, Y544^RBD^, Y557^RBD^, Y559^RBD^) and M460^RBD^. These residues form a pocket to wrap around the R328^ACE2^ helix with polar and stacking interactions (**Fig. 4c**). Patch 2 is mainly formed by β5 and its exiting loop, β6 and β7. Despite the 4.2 Å resolution, hydrogen bonds form between NAG386^ACE2^ and Y523^RBD^, A386^ACE2^ and Y521^RBD^, K352^ACE2^ and Y512^RBD^, N353^ACE2^ and E510^RBD^, N353^ACE2^ and Y544^RBD^. Y517^RBD^ on the tip of the β5-exiting loop makes extensive interactions with R26^ACE2^, D90^ACE2^, I92 ^ACE2^, and I93^ACE2^. A prominent feature of the ACE2 interface on RBD is that tyrosine-rich interactions provide 50% of the receptor-binding surface (**Supplementary Table 4**).

Comparison of the structures of ACE2-bound HKU5^RBD^, SARS-CoV-2^RBD^, PDF-2180^RBD^ and NeoCoV^RBD^ shows that HKU5^RBD^ binds to ACE2 in a different approaching mode (**Fig. 4d**). Both PDF-2180^RBD^ and NeoCoV^RBD^ bind to ACE2 mainly through their β5-exiting loops and the β-hairpins between strand β6 and β7 with smaller footprints. In contrast, HKU5^RBD^ and SARS-CoV-2^RBD^ recognize much larger areas on ACE2. While SARS-CoV-2^RBD^ grabs both sides of ACE2 α1 helix, HKU5 only interacts with the inner side of the α1 helix, shifting its footprint to the center region of ACE2 peptidase domain (**Fig. 4d**). Despite the common usage of ACE2 as receptor and similar folding of their RBDs, the subtle differences in the RBDs may led to distinct binding modes on ACE2.

### *P. abramus* ACE2 determinants of HKU5 spike interaction

To functionally validate residues in the binding interface, we generated both combinatorial and individual substitutions in *P. abramus* ACE2, transfected these constructs into BHK-21 cells, and used VSV-HKU5^spike^-RLuc or VSV-SARS2-CoV-2^spike^-RLuc to assess the ability of these constructs to support viral entry. We tested four combinations of substitutions: R26K/V30D/N38D/H41Y, D90N/I92T/I93V, P324Q/R328E/D329N, and N353G. Of these four ACE2 combinations, only P324Q/R328E/D329N (patch 1 in **Fig 4C**) completely abrogated HKU5 entry while N353G partially reduced entry (**Fig. 5a,b**). Each of individual substitution in P324Q/R328E/D329N also modestly reduced HKU5 entry.

**Fig. 5.**
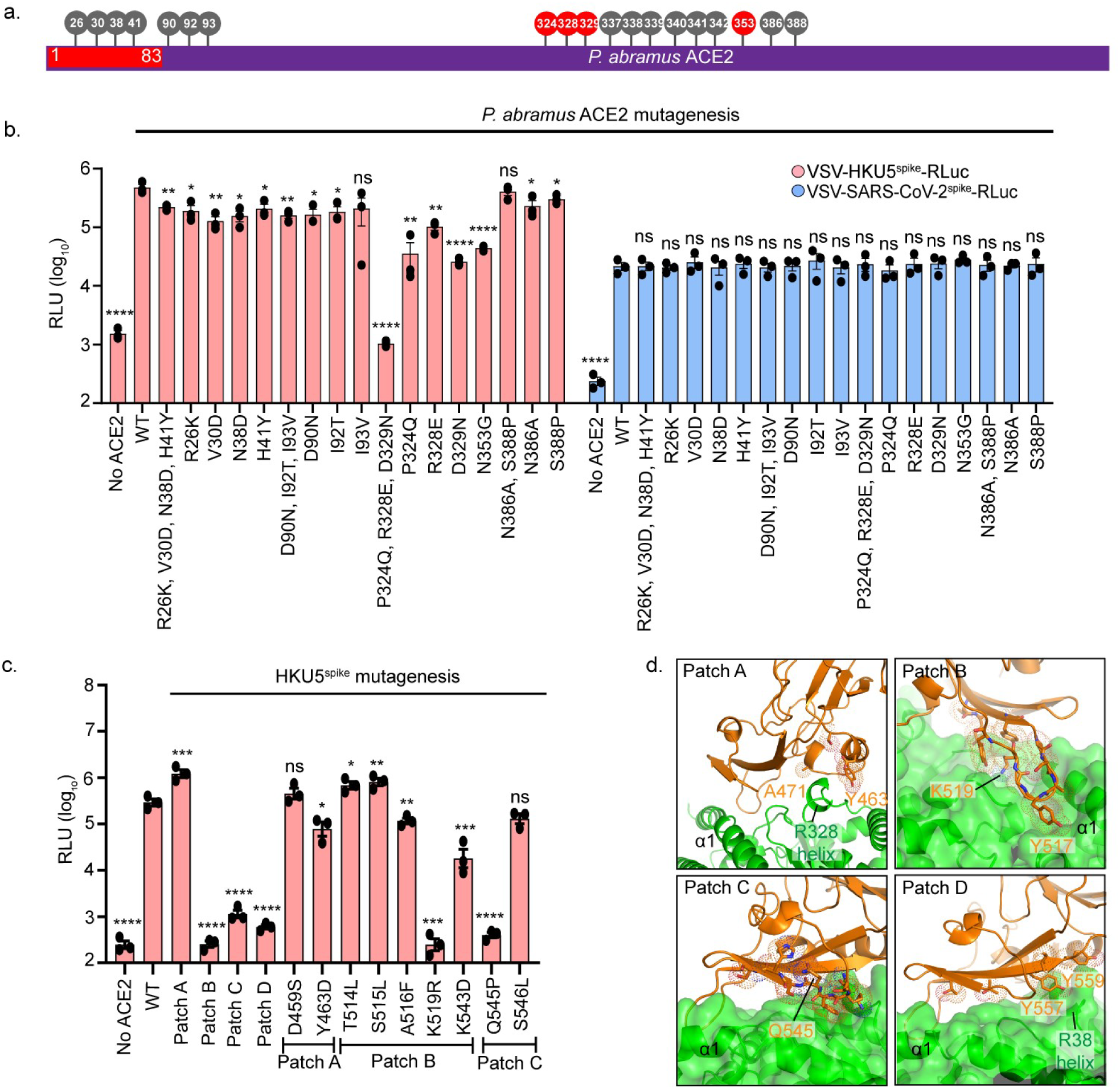
Molecular determinants affecting HKU5-ACE2-mediated entry. **a**, Schematic of the ACE2 gene highlighting substituted residues and domains. Positions in gray had minimal effect on HKU5 entry while substitutions that reduced entry are shown in red. **b,** Wild-type *P. abramus* ACE2, individual, and combinatorial ACE2 mutants were transiently transfected into BHK-21 cells and then infected with VSV-HKU5-spike-RLuc or VSV-SARS-CoV-spike-RLuc. **c,** We generated four patches of substitutions (A, B, C, D) on the HKU5^RBD^ and tested each for infectivity on BHK-21 cells stably expressing *P. abramus* ACE2 (Patch A:S457P/D459S/Y463D/A471P;patchB:T509N/E510K/Y512S/T514L/S515L/A516F/Y517D/G518D/K519R/ Y521E; patch C: 542-548◊EDGDYYRKQLSPLEG; patch D: T553A/T555S/Y557S/I558T/Y559V). HKU5 patches B, C, and D were required *for P. abramus* ACE2 use with individual substitutions K519R^RBD^ and Q545P^RBD^ sufficient to block infection. Dots represent the mean from each of three independent experiments each done in technical triplicate. Statistical analyses were performed using two-tailed unpaired Student’s *t*-tests and one-way ANOVA. Data are mean ± s.e.m. *\*P<0.05,**P<0.01, ***P<0.001,****P<0.0001.* (d) HKU5 RBD patches A, B, C and D. Residue positions for each set of substitutions are shown in stick and dot representation. K519^RBD^ and Q545^RBD^, which showed infection blocking effects when mutated, are labeled along with other residues for positional reference.

Although we did not previously observe detectable SARS-CoV-2 entry in *P. abramus* ACE2 expressing cells (**Fig. 1-2**), we hypothesized that a higher multiplicity of infection (MOI) using a concentrated VSV-SARS-CoV-2^spike^-RLuc stock could potentially overcome the barrier to infection. Indeed, our higher titer VSV-SARS-CoV-2^spike^-RLuc demonstrated viral entry with *P. abramus* ACE2 (**Fig. 5b**). Of note, none of the *P. abramus* ACE2 substitutions reduced SARS-CoV-2 entry, demonstrating that each ACE2 construct was functionally expressed on the cell surface.

An earlier study examining interactions between *Pipistrellus pipistrellus* ACE2 and NeoCoV/PDF-2180 revealed a glycan-dependent ACE2 binding mechanism, which mediates species-specific ACE2 use^28^. Specifically, *P. pipistrellus* residues 337-342^ACE2^ (SDGRQV) mediated entry of both NeoCoV and PDF-2180^28^. These residues are conserved in *P. abramus* ACE2 but differ between human and *P. abramus* ACE2 orthologs (Extended Data Fig. 6a,b). To determine whether these ACE2 determinants can modulate HKU5 entry, we generated six human ACE2-*P. abramus* ACE2 chimeras (Extended Data Fig. 7). We expressed the ACE2 chimeras in HEK-293T cells and validated expression by Western blot (Extended Data Fig. 7a, b). VSV-HKU5^spike^-RLuc and VSV-SARS-CoV-2^spike^-RLuc were used to assess the ability of these chimeras to support entry. Notably, introduction of the N-terminus of *P. abramus* ACE2 (1-83 aa, chimera #1) into human ACE2 enabled HKU5 entry; however, the reciprocal insertion of the human ACE2 N-terminus into *P. abramus* ACE2 (chimera #2) did not support entry (Extended Data Fig. 7a, b). Insertion of *P. abramus* residues 337-342^ACE2^ (SDGRQV) also did not facilitate HKU5 entry (chimera #3). Insertion of human residues 337-342^ACE2^ (GNVQKA, chimera #4) into *P. abramus* ACE2 modestly reduced but did not eliminate HKU5 entry. Next, we tested VSV-GD2013^spike^-RLuc to determine whether these interactions were generalizable to other HKU5-like viruses. VSV-GD2013^spike^-RLuc revealed a similar pattern to HKU5 amongst the ACE2 chimeras on HEK-293T cells (Extended Data Fig. 7c). We validated ACE2 chimera expression by Western blot (Extended Data Fig. 7d). In contrast to HKU5 and GD2013, SARS-CoV-2 was much more tolerant of ACE2 chimeras and substitutions. Both insertion of the *P. abramus* ACE2 N-terminus into human ACE2 (chimera #1), as well as the reciprocal insertion (chimera #2), were sufficient for SARS-CoV-2 entry (Extended Data Fig. 7c). Substitution of residues 337-342^ACE2^ had no effect on SARS-CoV-2 entry in cells expressing human or *P. abramus* ACE2, consistent with our structural analysis (Extended Data Fig. 7c).

To further investigate spike-ACE2 interactions, we conducted a flow cytometry-based binding assay with RBD-Fc fusion proteins on HEK-293T cells expressing the ACE2 chimeras. Consistent with the pseudovirus results, HKU5^RBD^ strongly bound both wild-type *P. abramus* ACE2 and the *P. abramus* ACE2 N-terminus on human ACE2 (chimera #1) (Extended Data Fig. 8a). Substitutions of residues 337-342^ACE2^ had no effect on HKU5^RBD^-Fc binding. In contrast to HKU5, SARS-CoV-2^RBD^-Fc bound to all ACE2 chimeras except chimera #4, phenocopying VSV-SARS-CoV-2^spike^-RLuc results. Next, we directly compared PDF-2180^RBD^-Fc binding relative to HKU5 with the ACE2 chimeras. PDF-2180 exhibited a broad but distinct pattern of ACE2 use relative to HKU5, consistent with different binding mechanisms and specificities (**Fig. 4d**). Consistent with previous results^28^, insertion of human ACE2 residues 337-342^ACE2^ (chimera #5) into *P. abramus* ACE2 impaired PDF-2180^RBD^-Fc binding (Extended Data Fig. 8c). Our structural analysis and mutagenesis results confirm that the N-terminus of *P. abramus* ACE2 is important for interaction with the HKU5^RBD^ and reveal distinct mechanisms of binding and entry between SARS-CoV-2, PDF-2180, and HKU5.

### HKU5 spike determinants of *P. abramus* and *N. vison* ACE2 interaction

We next sought to determine the HKU5^spike^ residues that interact with *P. abramus and N. vison* ACE2. We tested infectivity VSV-HKU5^spike^-RLuc in BHK-21 cells expressing either wild-type *P. abramus* ACE2 or *N. vison* ACE2. We introduced substitutions from MERS-CoV^spike^ on HKU5^spike^ at sites residing in the *P. abramus* ACE2 contact interface. HKU5^spike^ substitutions S457P/D459S/Y463D/A471P (Patch A) did not affect HKU5 entry in cells expressing *P. abramus* ACE2. Whereas HKU5^spike^ patch B (T509N/E510K/Y512S/T514L/S515L/A516F/Y517D/G518D/K519R/Y521E), patch C (542-548→EDGDYYRKQLSPLEG), and patch D (T553A/T555S/Y557S/I558T/Y559V) substitutions restricted infection with *P. abramus* ACE2. We further identified that the individual substitutions K519R (from patch B) and Q545P (from patch C) were sufficient to reduce *P. abramus* ACE2 use (**Fig. 5c, d**). In contrast, all four patches blocked infection of *N. vison* ACE2-expressing cells, with individual substitutions Y463D, S515L, A516F, K519R, K543D, Q545P, and S546L each being sufficient to restrict *N. vison* ACE2-dependent entry **(Extended Data 9)**. These findings suggest that *N. vison* ACE2 is more susceptible to HKU5^spike^ perturbations, which is consistent with the reduced HKU5 binding and entry efficiency relative to *P. abramus* ACE2.

### *N. vison* ACE2 determinants of HKU5 spike interaction

Our screen revealed that ACE2 from American mink (*N. vison*) but not ferrets (*Mustela putorius*) could support HKU5 entry (**Fig. 1**). Therefore, we asked which other mustelid species could serve as potential intermediate hosts for HKU5. To test this, we synthesized ACE2 constructs from nine additional mustelids including European badgers (*Meles meles*), greater hog badgers (*Arctonyx collaris*), Chinese ferret badger (*Meogale moschata*), North American river otters (*Lontra canadensis*), Eurasian otters (*Lutra lutra*), wolverines (*Gulo gulo*), stoats (*Mustela erminea*), black-footed ferrets (*Mustela nigripes*), and European mink (*Mustela lutreola*) (**Fig. 6a and Supplementary Table 2**). VSV-HKU5^spike^-RLuc and VSV-SARS-CoV-2^spike^-RLuc were used to assess the ability of wild-type mustelid ACE2s to support viral entry. *N. vison* and *M. erminea* ACE2 but not that of the other tested mustelids allowed for HKU5 entry (**Fig. 6b**). In contrast to VSV-HKU5^spike^-RLuc, VSV-SARS-CoV-2^spike^-RLuc similarly used all mustelid ACE2s (**Fig. 6b**). These findings suggest that *N. vison* and *M. erminea* could serve as intermediate host for HKU5-related viruses.

**Fig. 6.**
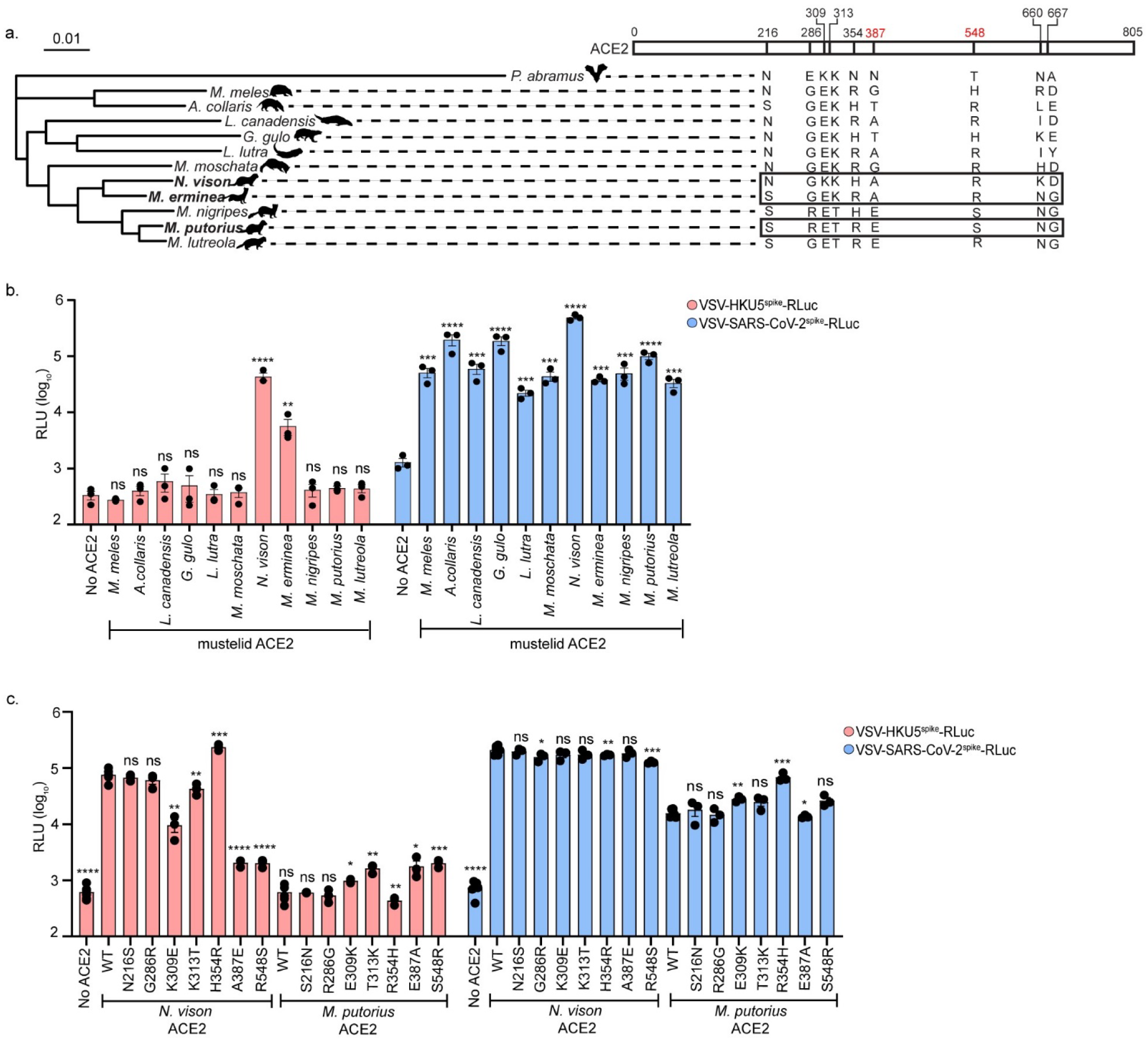
HKU5 interacts with mink ACE2 in a manner distinct from that of bat ACE2. **a**, Phylogenetic tree of mustelid species for which ACE2 constructs were tested. Shown are the nine amino acid positions that differ between *N. vison* (susceptible to HKU5) and *M. putorius* (resistant to HKU5) **b,** Wild-type mustelid ACE2 constructs were transiently transfected into BHK-21 cells and VSV-HKU5-^spike^-RLuc and VSV-SARS-CoV-^spike^-RLuc were used to assess entry. *N. vison* (American mink) and *M. erminea* (stoat) ACE2, but no the ACE2 of other mustelids, were sufficient for HKU5 entry. SARS-CoV-2 could use all tested mustelid ACE2s. **c,** Wild-type and mutant *N. vison* ACE2 and *M. putorius* ACE2s were transiently transfected into BHK-21 cells and entry was assessed using VSV-HKU5-^spike^-RLuc and VSV-SARS-CoV-^spike^-RLuc. *N. vison* ACE2 residues at A387^ACE2^ and R548^ACE2^ (highlighted in red) are necessary for HKU5 use of *N. vison* ACE2. No individual substitutions were sufficient for HKU5 use of *M. putorius* ACE2. Data are mean from three independent experiments each done in at triplicate. Statistical analyses were performed using two-tailed unpaired Student’s *t*-tests and one-way ANOVA. Data are mean ± s.e.m. *\*P<0.05,**P<0.01, ***P<0.001,****P<0.0001*.

To gain mechanistic insight into the *N. vison* ACE2 determinants of HKU5 entry, we leveraged the fact that *N. vison* and *M. putorius* have similar ACE2 sequences but dissimilar HKU5 susceptibilities. *N. vison* (susceptible) and *M. putorius* (resistant) differ at only nine sites across the ACE2 coding sequence (**Fig. 6a**). We induced reciprocal single amino acid substitutions at these sites in *N. vison* and *M. putorius* ACE2 in an attempt to modify the observed susceptibility of the wild-type ACE2 (**Fig. 6a**). Substitutions A387E and R548S on *N. vison* ACE2 reduced VSV-HKU5^spike^-RLuc entry, while no tested substitutions on *M. putorius* ACE2 fully restored HKU5 susceptibility (**Fig. 6c**). VSV-SARS-CoV-2^spike^-RLuc entered all constructs for both species, underscoring the broad ACE2 host range of SARS-CoV-2 (**Fig. 6c**). Position 387 represents an N-linked glycan site on *P. abramus* (N387^ACE2^) but not on that of *N. vison* (A387^ACE2^), *M. putorius* (E387^ACE2^), or human (A387^ACE2^). We identified that the glycan at N387^ACE2^ forms a hydrogen bond with Y523^RBD^; however, the role of 387^ACE2^ in mediating differences between mink and ferret ACE2 is independent of this glycan, as neither *N. vison* nor *M putorius* possess an N-linked glycan site at this position.

Our finding that 548^ACE2^ is required for HKU5 use of *N. vison* ACE2 was unexpected given this residue is remote from the binding interface, consistent with an allosteric effect. In addition, 548^ACE2^ was not identified as critical for HKU5 infection of *P. abramus* ACE2, PDF-2180 infection of *P. pipistrellus* ACE2, or SARS-CoV-2 infection of human or *N. vison* ACE2^9,39,40^ (**Fig. 6**). Position 548^ACE2^ was also not interrogated in previous ACE2 deep mutational scanning studies^41–43^. This position 548 represents the third residue in the NxS/T consensus sequence for a putative N-linked glycan site at 546^ACE2^ for *P. abramus* (T548^ACE2^), human (T548^ACE2^), and *M. putorius* (S548^ACE2^) but not *N. vison* (R548^ACE2^) nor *M. erminea* (R548^ACE2^) ACE2. Consistent with the distinct mechanism of interaction of HKU5-ACE2 relative to other coronaviruses, residues 337-342^ACE2^, which mediated PDF-2180 binding and entry, did not have an effect on *N. vison* ACE2 use by HKU5 (Extended Data Fig. 10). This suggests that differential ACE2 glycosylation plays an important role in mediating mustelid ACE2 use by HKU5 and highlights that HKU5 may interact with *N. vison* ACE2 in a distinct manner relative to *P. abramus* ACE2.

## DISCUSSION

Identifying viral receptors is critical for pandemic preparedness as receptor usage determines host range, cell and tissue tropism, pathogenesis, and potential for cross-species transmission^44^. Understanding virus-receptor interactions also informs the development of vaccines, therapeutic monoclonal antibodies, antiviral drugs, and diagnostics. In this study, we employed a diverse panel of receptor orthologs from 54 wild, domestic, and peri-domestic species to demonstrate that HKU5 efficiently utilizes ACE2 from *Pipistrellus abramus* (Japanese house bat), *Neogale vison* (American mink), and *Mustela erminea* (Eurasian ermine, stoat) as a receptor. This was confirmed using both pseudovirus and full-length HKU5 virus. Furthermore, we observed that the related merbecovirus GD2013 also engages *P. abramus* ACE2. Our findings that HKU5 uses *P. abramus* ACE2 was simultaneously confirmed by two additional groups^45,46^.

Cryo-EM analysis, coupled with mutagenesis studies, identified residues in *P. abramus* ACE2 that mediate HKU5^spike^ binding and entry. These residues are distinct from those involved in ACE2 interactions with SARS-CoV-2, PDF-2180, and NeoCoV, consistent with convergent evolution^28,29,47^. Notably, we observed that the critical ACE2 residues facilitating HKU5 entry in *N. vison* differ from those in *P. abramus*, revealing multiple, distinct mechanisms of ACE2 engagement among receptor orthologs. This observation further underscores the evolutionary plasticity of coronavirus spike-ACE2 interactions^48^.

Interestingly, HKU5 displayed a highly restricted ACE2 host range, successfully using only *P. abramus*, *N. vison*, and *M. erminea* ACE2 among the 57 orthologs tested. In contrast, other ACE2-using coronaviruses — including most sarbecoviruses (e.g., SARS-CoV and SARS-CoV-2), the alphacoronavirus HCoV-NL63, and the merbecoviruses PDF-2180 and NeoCoV—exhibit much broader ACE2 usage across mammalian species^18–21,28,49,50^. The narrow ACE2 host range of HKU5 is reflected in the distinct molecular interactions with *P. abramus* ACE2, differing substantially from those of SARS-CoV-2, PDF-2180, and NeoCoV. Importantly, sequence-based predictions alone would not have identified mink and stoat as potential intermediate hosts, as their ACE2 sequences do not phylogenetically cluster with *P. abramus*. This highlights the necessity of functional, unbiased experimental approaches for assessing coronavirus host susceptibility.

The narrow receptor usage of HKU5 raises an important evolutionary question as to why the HKU5 receptor host range is so constrained relative to other coronaviruses. One possibility is that HKU5 has evolved a highly specific adaptation to *P. abramus* ACE2, optimizing replication in its natural reservoir at the cost of reduced cross-species transmissibility. It is also possible that the co-existence of other coronaviruses in *P. abramus* bats may have induced antibodies that forced HKU5 to employ an alternative mode for binding ACE2. Given the evolutionary trajectory of other coronaviruses, HKU5 may acquire mutations that expand its ACE2 host range over time. Identifying the potential evolutionary pathways that could enable HKU5 to infect additional species—including humans—remains a critical area for future investigation.

*P. abramus* bats, commonly known as Japanese house bats, are ∼1.5-inch-long insectivorous bats abundant in East Asia that thrive in human-altered landscapes, often roosting in barns, bridges, and other man-made shelters. This places *P. abramus* bats in frequent contact with humans and farmed animals. While *N. vison* is native to only North America, they are routinely farmed and invasive in parts of Europe and Asia^51,52^ enabling interactions between *P. abramus* bats. Our findings here that HKU5 can use *P. abramus* and *N. vison* ACE2 have proven foreboding as during preparation of this manuscript, HKU5 was reported in two diseased minks on a Chinese mink farm^53^. This was the first description of HKU5 outside of *Pipistrellus* bats and was accurately predicted by our study. HKU5 infection of farmed American mink is of particular concern given multiple events of SARS-CoV-2 spillover from humans to mink and spillback to humans have been described highlighting mink as an effective intermediate host for coroanviruses^51,52,54,55^. The bunyavirus SFTSV which can cause hemorrhagic fever has also transmitted from mink to mink farmers further highlighting the zoonotic risk of farmed mink^56^.

Bats of the larger *Pipistrellus* genus, which are likely susceptible to HKU5, are distributed throughout Asia, Europe, and North Africa, thus presenting the opportunity for expanded HKU5 geographical range and further contacts with susceptible mustelids. It is possible that other mustelids beyond American mink and stoat will be susceptible to HKU5 viruses which may facilitate HKU5 evolution and host range expansion, perhaps even to humans. This underscores the need for increased surveillance of farmed and wild mustelids, particularly in areas with *Pipistrellus* bats^57,58^.

Our study identifies ACE2 as a previously unrecognized receptor for the merbecoviruses HKU5 and GD2013 and underscores mustelids, including mink and stoats, as potential intermediate hosts. Group 2c merbecoviruses including HKU5 have pandemic potential and should be considered in the design of merbecovirus vaccines. Our findings demonstrate that SARS-CoV-2 and MERS-CoV vaccine sera poorly neutralize HKU5, underscoring the need for pan-merbecovirus vaccine strategies. The identification of *P. abramus* ACE2 as a receptor for HKU5 will facilitate these efforts. These findings have important implications for zoonotic spillover risk and underscore the urgent need for continued surveillance of coronaviruses in both their natural reservoirs and potential bridging hosts.

## MATERIALS AND METHODS

### Receptor and virus sequences

The sequence information of the 31 mammalian DPP4s and 54 mammalian ACE2s, directly retrieved from GenBank, is described in Supplementary Table 1 and Supplementary Table 2. The whole genome sequences of selected coronaviruses were retrieved the following GenBank accessions: SARS-CoV2 (MT461669), bat HKU5-LMH03f (NC_009020.1)^59^, PDF-2180 (NC_034440.1)^60^, NeoCoV (KC869678.4)^61^, bat Hp-BetaCoV/Zheijiang2013 (NC_025217.1)^62^, RoBatCoV HKU9 (NC_009021.1)^25^, RoBatCoV GCCDC1 (NC_030886.1)^63^, *Erinaceus* CoV/2012-174 (NC_039207.1)^64^, *Erinaceus* CoV/HKU31strainF6 (MK907286.1)^65^, BatCoV HKU5 isolate BY140568 (MN611520.1)^33^, BtPa-BetaCoV/GD2013 (KJ473820.1)^34^, BatCoV HK25 isolate NL140462 (KX442565.1), BatCoV HKU25 isolate YD131305 (KX442564.1), BtVs-BetaCoV/SC2013 (KJ473821.1)^35^, BtCoV/133/2005 (Q0Q4F2.1)^66^, BtTP-BetaCoV/GX2012 ( KJ473822.1)^34^, BatCoV/H. savii/Italy/206645-40/2011 (MG596802.1)^36^, BatCoV HKU4r isolate GZ131656 ( MN611519.1)^33^, BatCoV HKU4 (NC_009019.1)^17^, BatCoV HKU4r isolate SM3A (MW218395.1), Bat-SL-CoVZC45 (MG772933.1)^67^, human NL63 (JX504050.1)^68^, human 229E (MT797634.1), camel MERS KFU-HKU19Dam (KJ650296.1)^69^, and human MERS-CoV (NC_019843.3)^6^. Genes synthesized in this study were produced by GenScript or Twist.

### Plasmids

Human codon-optimized sequences of DPP4s, ACE2s, ACE2 chimeras, and ACE2 mutants with a C-terminal HA-tag were commercially synthesized and subcloned into an expression vector (pUC57) through the BsmbI restriction site. The DNA sequences of human codon-optimized coronavirus spikes and HKU5 mutants with a C-terminal Flag tag were commercially synthesized and subcloned into the pCAGGS vector through the EcoRI and NotI restriction sites. Plasmids expressing coronavirus spike N-terminal domain (NTD)-IgG-mFc and receptor binding domain (RBD)-IgG-mFc fusion proteins were commercially synthesized and generated by inserting the putative coding sequence of HKU5^NTD^ (amino acids 22-359), SARS-CoV-2^RBD^ (amino acids 316-527), HKU5^RBD^ (amino acids 376-586), MERS-CoV^RBD^ (amino acids 368-588), and PDF-2180^RBD^ (amino acids 372-584) into a modified pFuse-mIgG1-Fc2 vector (InvivoGen) through the EcoRI and NheI restriction sites. An IL2 signal sequence was included for protein secretion.

### Cells lines

HEK-293T and BHK-21 cells were cultured in Dulbecco’s Modified Eagle Medium (DMEM, Gibco) with 10% heat-inactivated fetal bovine serum (FBS) and 1% Penicillin/Streptomycin (Pen/Strep). Vero-CCL-81 cells were cultured in DMEM containing 5% FBS, 1% Pen/Strep, and 1% NEAA. For the Vero-CCL-81 ACE2 stable cell lines, 10 µg/ml of puromycin (Gibco) was added. Cells were cultured at 37°C in 5% CO2 with regular passaging every 2-3 days. Expi293F cells used for protein production was cultured in Expi293 Expression Medium (ThermoFisher Scientific), according to the manufacturer’s instructions. Expi293F cells were maintained at 37°C at 8% CO2 in an orbital shaker (125 rpm) and split at 0.3-0.5 **×** 10^6^ viable cells/ml every 3-4 days. BHK-21 cells were stably transduced with a lentiviral vector encoding *P. abramus* ACE2 for pseudovirus experiments with HKU5 mutants.

### Production of VSV-ΔG pseudoviruses

Vesicular Stomatitis Virus (VSV)-based pseudotyped viruses were produced as described previously^70–73^. pCAGGS containing different coronavirus spikes were synthesized by GenScript. Briefly, HEK-293T cells were transfected with pCAGGS vector expressing the different spikes using polyethylenimine (PEI) and after 24 h, were infected with a replication-deficient VSV vector containing expression cassettes for Renilla Luciferase (RLuc) or eGFP in lieu of the VSV-G open reading frame. After an infection period of 1 h at 37°C, the inoculum was removed, and cells were washed with 1X PBS prior to the addition of media supplemented with anti-VSV-G clone I4 (Kerafast) to neutralize residual input virus^74^. Pseudotyped particles were harvested at 24 and 48 hpi, clarified from cellular debris by centrifugation, and stored at −80°C prior to use.

### Production and purification of spike-RBD and NTD proteins

RBD- and NTD-mFc fusion proteins (SARS-CoV-2^RBD^ (amino acids 316-527), HKU5^RBD^ (amino acids 376-586), MERS-CoV^RBD^ (amino acids 368-588), PDF-2180^RBD^ (amino acids 372-584), and HKU5^NTD^ (amino acids 22-359) were expressed in Expi293F cells by transfecting the corresponding plasmids using the Expi293 expression system kit (ThermoFisher Scientific) following the manufacturer’s instructions. Briefly, Expi293F cells were subcultured in 125-ml shaker flasks. Once the cell density reached 2.5 **×** 10^6^ viable cells/ml in a 45 ml culture, 45 µg of plasmids were transfected using the Expifectamine 293 transfection kit. After 20 hours post transfection (hpt), Enhancer 1 and 2 from the kit were added and transfected cells were incubated for an additional 4 days in an orbital shaker at 125 rpm at 37°C with humidified 8% CO2. The supernatant was subsequently harvested by centrifugation at 2000 rpm for 2 min. The protein-containing supernatant was transferred to a sterile 50-ml Falcon tube and mixed with pre-washed Protein A agarose resin in 1X PBS (GoldBio). The mixture was incubated at room temperature with gentle rocking for 2 h. The protein-bound agarose resin was purified through a gravity flow column (BioRad) and washed with 1X PBS. Before storing at −80°C, RBD proteins were eluted with 100 mM glycine (pH 3.2) and 5M NaCl and neutralized by 100 mM Tris-HCl (pH 8). The purified RBD and NTD proteins were visualized on SDS-PAGE gel by Coomassie Blue staining using the BioRad ChemiDoc Imaging System.

### Pseudotyped virus entry assay

HEK-293T or BHK-21 cells were seeded in 100 µl total volume in each well of a black-walled clear bottom 96-well plate. The following day, cells were transfected with either DPP4- or ACE2-expressing plasmids using PEI. The following day, spike expressing VSV-pseudotype viruses was added at 1:10 final concentration volume/volume (v/v) in DMEM with 2% FBS and incubated for 24 h. After 24 h, cells were lysed, and luciferase activity was quantified. In brief, 5X Renilla lysis buffer (Promega) was diluted using 1X PBS. 25 µl of 1X lysis buffer was added to each well prior to incubating for 20 min at room temperature (shaking). Renilla Luciferase substrate (Promega) was mixed with Renilla assay buffer (10 µL of Renilla Luciferase substrate per ml of assay buffer) and added to the lysed cells at a volume of 25 µl. The cells were incubated for 10 min on a plate shaker in the dark. Plate measurements were taken in duplicate with biological replicates using a microplate reader (BioTek Synergy) and Gen5 software and the relative light units (RLU) were plotted and normalized in GraphPad Prism (v10.2)

### MLV-ACE2 entry into spike-expressing cells

Murine leukemia virus luciferase reporter virions bearing *P. abramus* ACE2 (VSV-*P. abramus*^ACE2^-Gluc) were tested for entry into spike expressing HEK-293T cells. MLV-ACE2 virions were produced by transfecting 0.4 µg of MLV-GagPol plasmid (pMDoldGag-Pol, Richard Mulligan, Harvard Medical School, Boston, MA, USA), 0.4 µg of ACE2 plasmid or empty vector, and 0.2 µg of an intron-regulated MLV-based Gaussia Luciferase (GLuc) reporter gene (Dave Derse, NCI, Frederick, MD, USA) into a 24-well plate of HEK-293T cells using PEI^75^. VSV-*P. abramus*^ACE2^-GLuc containing supernatants were purified with a 0.45 µm syringe filter two days post transfection (dpt) and used fresh. HEK-293T cells were 0.2 µg spike plasmids or empty vector using PEI into a 48-well plate 1 day prior to the assay. Purified VSV-*P. abramus*^ACE2^-Gluc supernatants (100 µl) were added to spike-expressing target cells in quadruplicate and incubated at 37°C. After two days, 100 µl of assay supernatant were mixed with 30 µl Gluc substrate (ThermoFisher Scientific) containing 100 mM NaI in a white 96-well plate and subsequently read for luminescence with a Promega GloMax Explorer plate reader.

### Spike RBD-Fc and NTD-Fc protein binding assay

HEK-293T cells transiently transfected with *P. abramus* ACE2 were trypsinized and incubated with 50 µg/ml of recombinant RBD- or NTD-mFc protein diluted in DMEM with 2% FBS. The cell and Fc-protein mix was incubated for 2 h in a roller at 4°C (cold room). Unbound spike RBD- or NTD-mFc proteins were removed and cells were washed with cold 1X PBS. For flow cytometry analysis, spike RBD- or NTD-mFc bound cells were incubated with a live/dead stain (Zombie Violet, Biolegend) in 1X PBS for 10 min at 4°C and fixed in 4% paraformaldehyde (PFA) for 10 min at room temperature. Following cell fixation, 100 µl (1:500) of Alexa Fluor 488-conjugated goat anti-mouse IgG (Invitrogen) was added into the spike RBD-mFc or NTD-fc bound cells and incubated for 30 min at 4°C. The cells were then washed and resuspended with cold FACS buffer (1X PBS, 2% FBS, 1 mM EDTA). Dead cells, as indicated by SSC/FSC, were excluded by gating. The spike RBD-mFc or NTD-Fc bound cells were assessed by the fluorescence intensity compared to HEK-293T control cells without the receptor expression. Flow cytometry data were collected on a Beckman Coulter CytoFlex S and analyzed using FlowJo (v.10).

### Infection with authentic full-length HKU5 virus

*P. abramus* and *N. vison* ACE2-expressing cells were infected with full length-HKU5 infectious clone virus (FL-HKU5, multiplicity of infection (MOI): 0.01) for 72 h. CPE was monitored and images were taken using an inverted optical microscope (Olympus, IX73). For the multistep growth curve analysis, cells were infected at a MOI of 0.01 and incubated at 37°C with 5% CO2 for 1 h. After removing the inoculum, the monolayer was washed with 1X PBS. Complete growth media was added back to each well and samples of the infected culture supernatant were collected at 0, 4, 24, 48, and 72 hpi. Samples were stored at −80°C for viral titration, which was performed by plaque assay on Vero81 cells expressing *P. abramus* ACE2 or *N. vison* ACE2 (Vero81-PaACE2 and -NvACE2, respectively). 10-fold serial dilutions of virus in 1X PBS were prepared and used to inoculate Vero81-PaACE2 and -NvACE2 cells, as described above. After 1 h of viral absorption, the monolayers were overlaid with 0.8% agarose in Eagle minimum essential medium (MEM). Plaques were visualized and manually quantified at 4 days post infection (dpi) using neutral red stain.

### Western blot

HEK-293T cells transfected with ACE2 or coronavirus spikes were washed with 1X PBS and lysed with NP-40 lysis buffer (ThermoFisher Scientific) on ice for 10 min. Cell lysates were clarified by centrifugation at 10,000 **×** g at 4°C for 5 min. The lysate was mixed with 4X Laemmli sample buffer (BioRad) at a 1:4 (v/v) ratio of buffer to lysate and incubated at 95°C for 5 min. For HEK-293T cells transfected with DPP4 orthologs, cells were lysed using RIPA buffer on ice for 10 min, mixed with 4X Laemmli sample buffer (BioRad) at a 1:4 (v/v) ratio of buffer to lysate, and loaded without boiling. To detect the Flag-tag on pseudotyped viruses, the virus-containing supernatant was mixed with 4X Laemmli sample buffer. After SDS-PAGE and PVDF membrane transfer, the blots were blocked with 5% milk in 1X TBS containing 0.1% Tween-20 (TBST) at room temperature for 1 h. Primary antibodies against Flag (Sigma-Aldrich), HA (Biolegend), and GAPDH (Biolegend) were added at 1:1000 dilution in TBST with 5% milk and incubated on a shaker at 4°C overnight. After three washes with TBST, the blots were incubated with horseradish peroxidase (HRP) conjugated secondary antibody goat anti-mouse IgG (H+L) (Jackson Immuno Research, 1:5000 dilution) in 1X TBST for 1 h on a shaker at room temperature. The blots were washed three times with 1X TBST and visualized using the BioRad ChemiDoc Imaging system.

A full-length molecular clone for HKU5 (FL-HKU5) has been previously reported^26,27^. To detect the nucleocapsid in FL-HKU5 virus infection, HEK-293T cells were transiently transfected with ACE2. Following 24 hpt, the cells were exposed to FL-HKU5 (MOI: 1.0), and the infected cells were collected at 24 hpi. These cells were then lysed using 8M urea, combined with Laemmli sample buffer (BioRad), and heated at 60°C for 10 min. After SDS-PAGE and PVDF membrane transfer, the blots were blocked with 5% milk in 1X PBS at room temperature for 1 h. Primary antibodies targeting spike (1:2000, gift from David Veesler, University of Washington, clone 76E1)^76^, HA (1:2000, CST catalog number: 3724), nucleocapsid (1:2000, Sino Biological 40068-RP02), and GAPDH (1:5000, CST catalog number: 2118) were diluted in PBS with 5% BSA and 0.1% tween 20 and left to incubate on a shaker at 4°C overnight. Following three washes with 1X PBS, blots were exposed to horseradish peroxidase (HRP) conjugated secondary antibodies goat anti-human (1:5,000, SeraCare catalog number: 5220-0330) and goat anti-rabbit (1:5,000, CST catalog number: 7074S) in 1X PBS with 5% BSA and 0.1% tween 20 for 1 h on a shaker at room temperature. Finally, the blots were washed three times with 1X PBS and visualized using the iBright 1500 Imaging system (Invitrogen).

### Immunofluorescence assay

For pseudovirus experiments, HEK-293T and BHK-21 cells were plated in 8-well chamber slides and transfected with ACE2, infected with pseudotyped-Spike-eGFP virus at 1:10 final concentration v/v in DMEM with 2% FBS, and incubated for 24 h. Cells were subsequently fixed with 4% PFA at room temperature for 10 min. The nuclei were stained with Hoechst 33342 (1:10,000 dilution in 1X PBS, ThermoFisher Scientific). Images were captured with a fluorescence microscope (Zeiss Axio Imager) and analyzed using the Zeiss Zen Pro microscopy software.

For FL-HKU5 experiments, Vero81-PaACE2 cells were seeded on coverslips in 6-well plates and subsequently infected with FL-HKU5 virus (MOI: 0.01). At 24 hpi, cells were fixed with 4% PFA for 15 min at room temperature. Cells were then blocked and permeabilized in 10% normal goat serum and 0.3% triton-x 100 for 1 h at room temperature, followed by incubation with rabbit anti-MERS polyclonal sera (1:2000; Sino Biological; 40068-RP02) diluted in 5% normal goat sera for 16 hours at 4°C. Cells were then washed three times with 1X PBS and incubated with goat anti-rabbit conjugated to Alex488 (1:3000) diluted in 5% normal goat sera at room temperature for 2 h. Nuclei were stained with Hoechst (1:10,000) and the cells were washed three times prior to mounting on slides with ProLong Gold antifade reagent (Invitrogen). HKU5 antigen-positive cells were observed under a fluorescence microscope (Keyence, BZ-X810) and images were captured for further analysis.

### Bioinformatic and computational analyses

Protein sequence alignment was performed using the MUSCLE algorithm by MEGA-X software (v.10.17)^77,78^. For phylogenetic analysis, amino acid sequences of receptors or spikes were first aligned using MUSCLE and phylogenetic trees were generated using the maximal-likelihood method in MEGA-X (1,000 boostraps). The model and parameters utilized for the phylogenetic analysis were implemented based on the recommended optimal protein model using the MEGA-X software. SimPlot (v.3.5.1) was used to analyzed nucleotide similarities with a sliding window size of 200 nucleotides and a step size of 20 nucleotide using a gap-stripped alignments and Kimura (2-parameter) distance model.

### Cryo-EM grid preparation, data collection and structure determination

The complex of *P. abramus* ACE2-ACT66266.1 and HKU5-RBD-10lnQQAVI was assembled by mixing ACE2 and HKU5-RBD at a molar ratio of 1:1.2 to a final concentration of the complex of 3.7 mg/ml, followed by incubating for 1 h at room temperature. Before vitrification, 6mM 3-[(3-cholamidopropyl) dimethylammonio]-2-hydroxy-1-propanesulfonate (CHAPSO) was added to the sample. A sample volume of 2.8 µl was applied onto a freshly glow discharged Quantifoil R 2/2 gold grid and plunge-frozen using a Thermo Scientific Vitrobot Mark IV plunger with the following parameters: chamber humidity of 95%, chamber temperature of 4°C, blotting force of −5 and blotting time between 1 s and 2.5 s.

The data were acquired using SerialEM^79^ on an FEI Titan Krios G1 electron microscope equipped with a Direct Electron Apollo direct electron detector. A total of 10,702 movies were collected at a nominal magnification of 47,000x with the pixel size of 0.5 Å in the super-resolution mode. Movie frames were aligned with MotionCor2^80^, and CTF parameters were estimated using cryoSPARC 4.4^81^ module patch CTF estimation. 9,529 micrographs were selected for downstream processing after curating in cryoSPARC based on full-frame motion, CTF fit resolution and relative ice thickness. Particles were picked from a subset of 500 micrographs using crYOLO^82^. After several rounds of 2D classification, selected 2D templates and corresponding particles were used for template picking and Topaz training^83^. The particles obtained by template-based and Topaz picking were merged, followed by elimination of duplicates. *Ab initio* reconstruction in cryoSPARC was used to generate initial models, followed by multiple rounds of heterogeneous refinement to remove low-quality particles. The resulting 242,675 particles representing the complex dimer were subjected to non-uniform refinement in cryoSPARC with C2 symmetry imposed to produce the final map of ACE2-HKU5RBD. The reported resolution was determined based on the “gold standard” criterion at the FSC curve threshold of 0.143^84^.

To build the atomic model of the complex, an initial model of *P. abramus* ACE2-ACT66266.1 was generated using SWISS-MODEL^85^ based on the structure from PDB entry 8WBY, while the initial model of HKU5-RBD-10lnQQAVI was obtained using ColabFold^86^. These models were docked into the cryo-EM map using UCSF Chimera^87^ and refined in Coot^88^ using the sharpened map generated by cryoSPARC and map post-processed using EMReady^89^. The improved model was refined using real-space refinement in Phenix^90^ against the original cryoSPARC map. Molprobity^91^ was used to validate the final model. The refinement statistics are summarized in the Supplementary Table 3.

### Production and purification of soluble ACE2 ectodomain proteins and biotinylated RBD proteins

N-terminus HRV3C cleavable single-chain Fc-tagged *P. abramus* ACE2 ectodomain proteins, and N-terminus HRV3C cleavable single-chain Fc-tagged and C-terminus AVI-tagged HKU5 RBD and SARS-CoV-2 RBD proteins were expressed in Expi293F cells by transfecting the corresponding plasmids using the Turbo293 transfection reagent (Speed BioSystems). Briefly, pre-mixed 1 mg plasmid in 20 ml Opti-MEM (Thermo Fisher Scientific) and 3 ml of Turbo293 transfection reagent in 20 ml Opti-MEM were added to 0.8 liter of Expi293F cells at cell density of 2.5 **×** 10^6^ viable cells/ml. The transfected cells were incubated for 6 days in an orbital shaker at 125 rpm at 37°C in a humidified 9% CO2 incubator before the supernatant was harvested by centrifugation and filtration. Subsequently, the supernatant was incubated with 5 ml of PBS-equilibrated protein A resin for 1-2 h. The ACE2-bound resin was then collected, washed with 1X PBS, and cleaved overnight at 4°C with 200 μg of HRV3C. The RBD-bound resin was collected, washed with 1X PBS, and subjected to an overnight incubation in a 3-ml BirA biotin-protein ligase mixture (Avidity) and 200 μg of HRV3C at 4°C. The liberated soluble ACE2 proteins and biotinylated RBD proteins were applied to a Superdex 200 16/600 gel filtration column equilibrated with PBS the next day. Peak fractions corresponding to the target proteins were pooled for subsequent analysis.

### Binding analysis using biolayer interferometry (BLI)

Binding affinities were assessed using BLI assays conducted on the Octet HTK instrument. Briefly, biotinylated recombinant RBD proteins at a concentration of 3 µg/ml were immobilized on streptavidin biosensors for 300 s. Following this, the biosensors were immersed in kinetic buffer for 180 s to remove any unbound biotinylated RBD or NTD proteins. Subsequently, the biosensors were immersed in kinetic buffer containing soluble *P. abramus* ACE2 ectodomain proteins at concentrations ranging from 0-500 nM for 120 s to capture association kinetics and then placed in kinetic buffer for 300 s to capture dissociation kinetics. A kinetic buffer lacking ACE2 was used as a reference for background determination. The affinities were then calculated Octet Data Analysis software (v.12.2.0.20), employing curve-fitting kinetic analysis or steady-state analysis with global fitting. Kd, app values were reported due to the utilization of dimeric ACE2.

### Neutralization assay

For pseudotyped virus neutralization, BHK-21 cells were transfected with plasmids containing the full-length human ACE2, human DPP4, and *P. abramus* ACE2 using PEI. The following day, two-fold serial dilutions of sera and antibody were prepared in DMEM. SARS-CoV-2 (Moderna) vaccine sera was pooled from healthy humans (BEI Cat# NRH-21747). MERS-CoV-27 monoclonal antibody was described previously^36,37^. MERS-CoV-S2P serum was harvested (Day 60) from mice primed (Day 0) and boosted (Day 30) with MERS-CoV-S2P (full length spike) and pooled from 10 mice. Subsequently, 5 µl of the corresponding pseudovirus was mixed with 20 µl of DMEM with 2% FBS and 25 µl of each serum/antibody dilution, and the mixtures were incubated for 45 min at 37°C. Transfected BHK-21 were trypsinized and seeded into 96-well plates at a density of 50,000 cells per well with the pseudovirus-sera/antibody mixtures. After 24 hr, the cells were lysed and Renilla luciferase activity was measured as described above.

### Statistical analysis

Infection and binding assay experiments were repeated at least 2-3 independent times with 2-4 technical repeats per experiment. Data are presented as mean ± s.e.m as specified in the figure legends. Statistical analyses were conducted using GraphPad Prism (v10.2) using unpaired two-tailed Student’s tests or One-way ANOVA, as stated in the figure legends. *P< 0.05* was considered significant; \**P<0.05*, \*\**P< 0.01*, *\*\*\* P<0.001*, \*\*\*\**P<0.0001*.

### Data availability

The structural data generated in this study are available in the Protein Data Bank (PDB) under accession number 9MV0 and in the Electron Microscopy Data Bank (EMDB) with an entry ID EMD-48650.

## Acknowledgements

This project has been funded in whole or in part with Federal funds from the National Cancer Institute, National Institutes of Health, under Prime Contract No. 75N91019D00024, Task Order No. 75N91023F00016. The content of this publication does not necessarily reflect the views or policies of the Department of Health and Human Services, nor does mention of trade names, commercial products or organizations imply endorsement by the U.S. Government. This work was funded in part by Burroughs Wellcome Fund (CBW), NIH R01 AI148467 (CBW), NIH R21 AI173821 (SL and CBW), NIH PO1 AI167966 and CEIRS Contract 75N93021C00014 (RSB), R01 AI183155 (SL), NIH grant R01 AI163395 (WM), Hanna H. Gray Fellowship from the Howard Hughes Medical Institute (DRM), NIH grant F31 AI176650 (MWG), and Metropolitan AntiViral Drug Accelerator (CBW). BLM was supported by NIH T32 HL007974. AS was supported by the Fullbright Junior Research Award Scholarship, PL/2023/21/JR. MWG is a recipient of Gruber Science Fellowship and was supported by NIH T32 AI055403. We thank Peter Cresswell, Matthew Spencer, and Leo Phan for helpful discussions and technical guidance.

---

**Extended Data Fig. 1.**
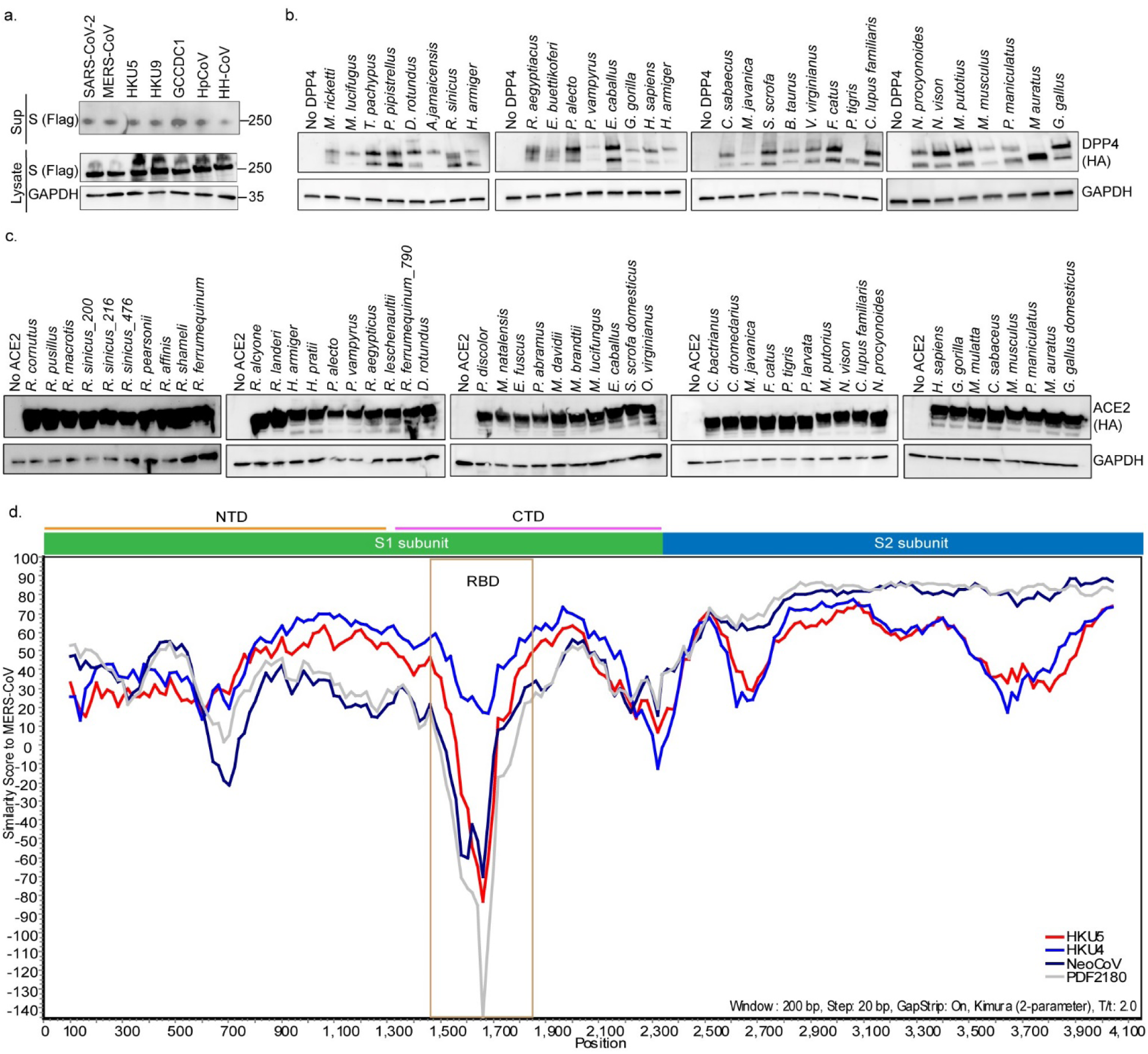
Western blot analysis of spikes and receptors. **a**, Western blot detected the expression levels of coronavirus spikes on HEK-293T cells by targeting the C-terminal Flag-tag. Sup: supernatant. **b,** Expression levels of dipeptidyl peptidase-4 (DPP4) orthologs transiently transfected in HEK-293T cells were detected by Western blot by targeting the C-terminal HA-tag. **c,** Expression levels of angiotensin converting enzyme 2 (ACE2) orthologs transiently transfected in HEK-293T cells were detected by Western blot by targeting the C-terminal HA-tag. Glyceraldehyde 3-phosphate dehydrogenase (GAPDH) was employed as a loading control. **d,** SimPlot analysis showing the spike nucleotide similarity of four merbecoviruses compared to MERS-CoV.

**Extended Data Fig. 2.**
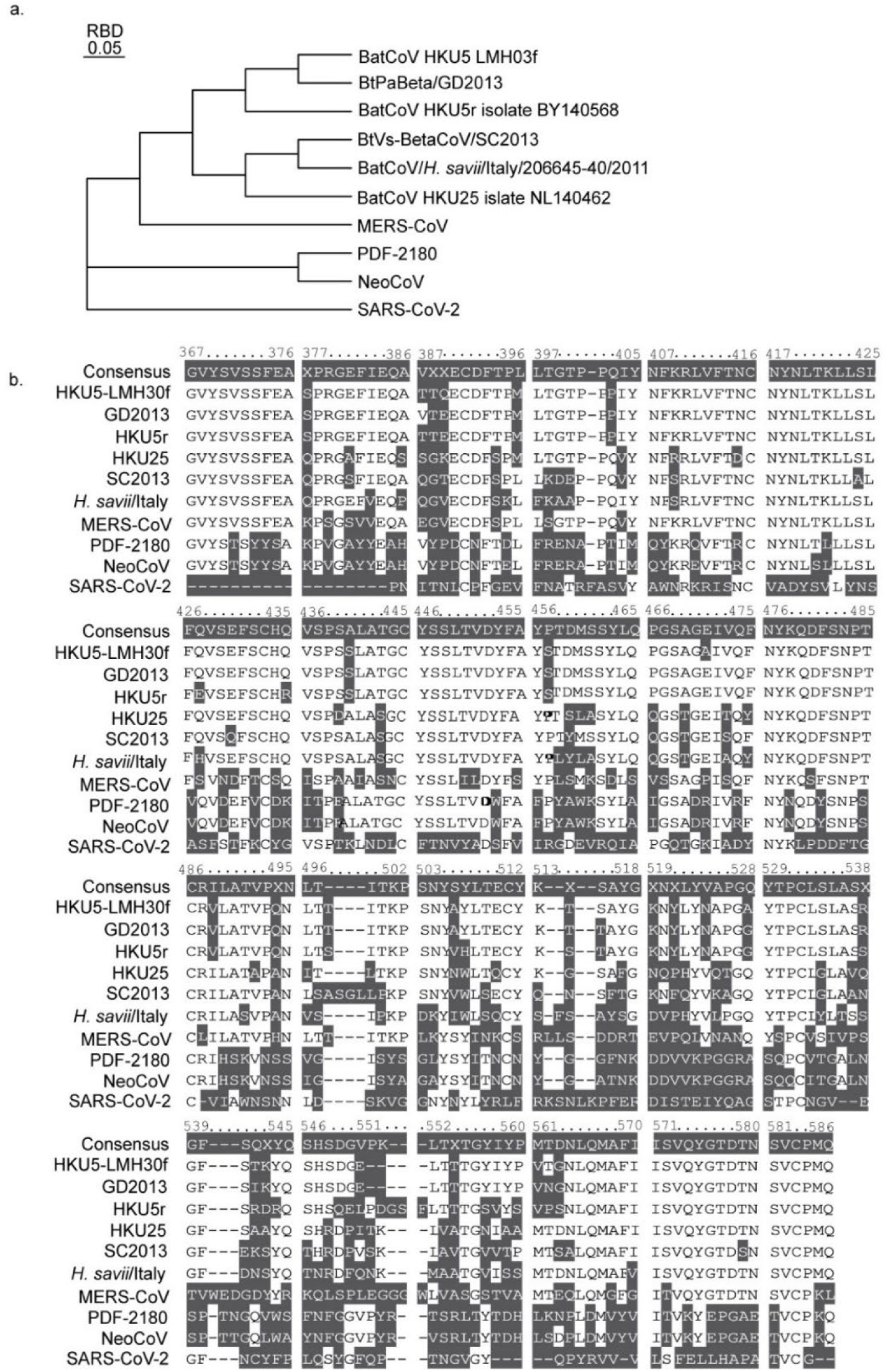
Phylogenetic tree and alignment of beta-coronavirus receptor binding domains (RBDs). **a,** HKU5 is closely related to GD2013 which can also use *P. abramus* ACE2 as a receptor. **b,** MUSCLE alignment of HKU5-like viruses highlights variable RBD sequences.

**Extended Data Fig. 3.**
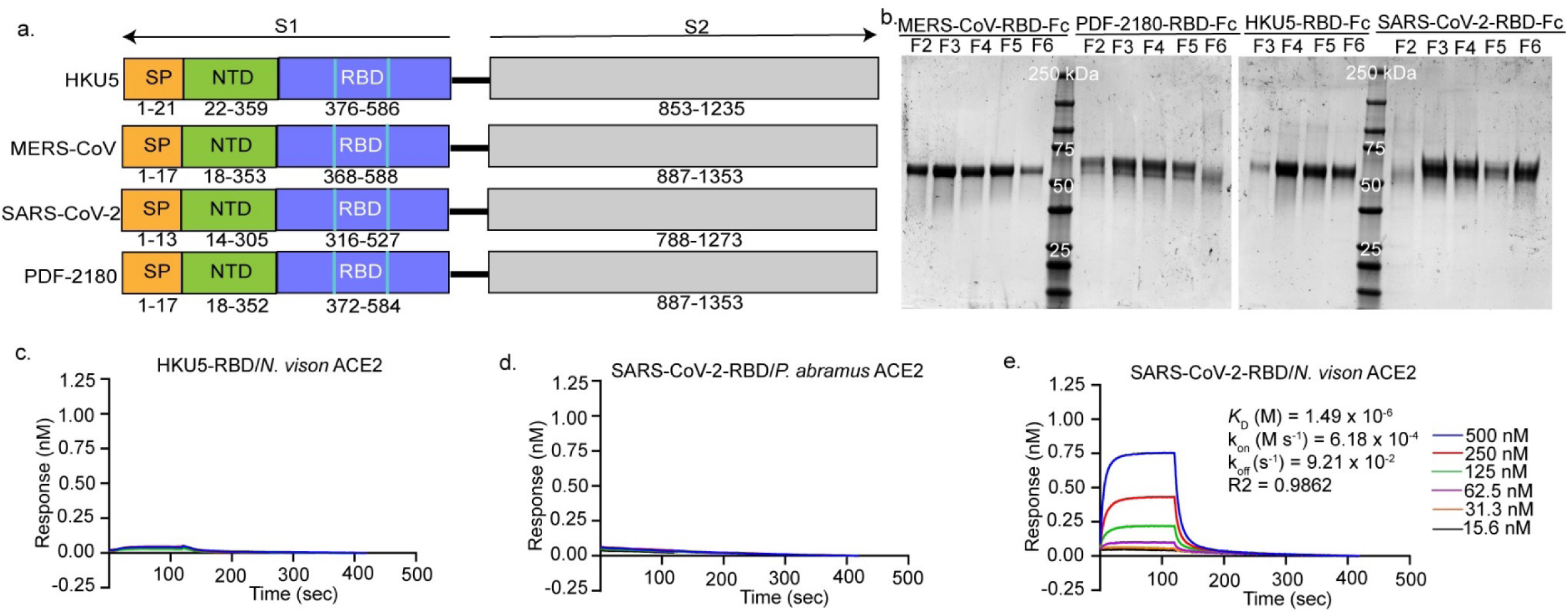
HKU5 RBD does not interact with *N. vison* ACE2. **a**, Schematic representation of spike protein of HKU5, MERS-CoV, SARS-CoV-2, and PDF2180. SP: Signal peptide; RBD: receptor binding domain; NTD: N-terminal domain. **b,** Purified spike RBD proteins were loaded on SDS-PAGE gels and stained with Coomasie blue to visualize the proteins. **c-e**, BLI assays analysis binding kinetics between HKU5-RBD with N. vison ACE2 (**c**), SARS-CoV-2-RBD with *P. abramus* bat ACE2 (**d**), and SARS-CoV-2-RBD with *N. vison* ACE2 (**e**).

**Extended Data Fig. 4.**
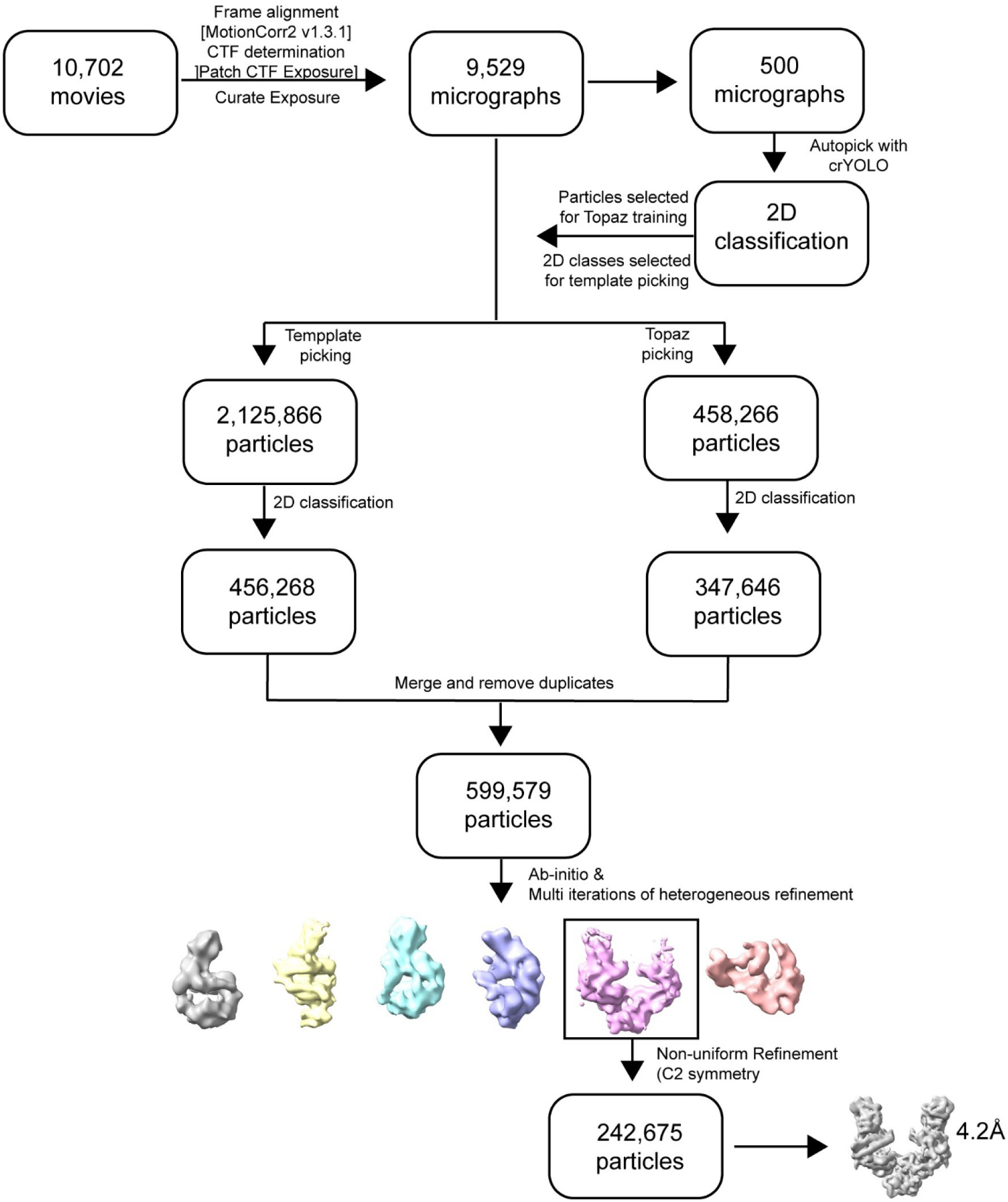
Cryo-EM data processing workflow for the complex of *P. abramus* ACE2 and HKU5-cytoplasmic tail domain.

**Extended Data Fig. 5.**
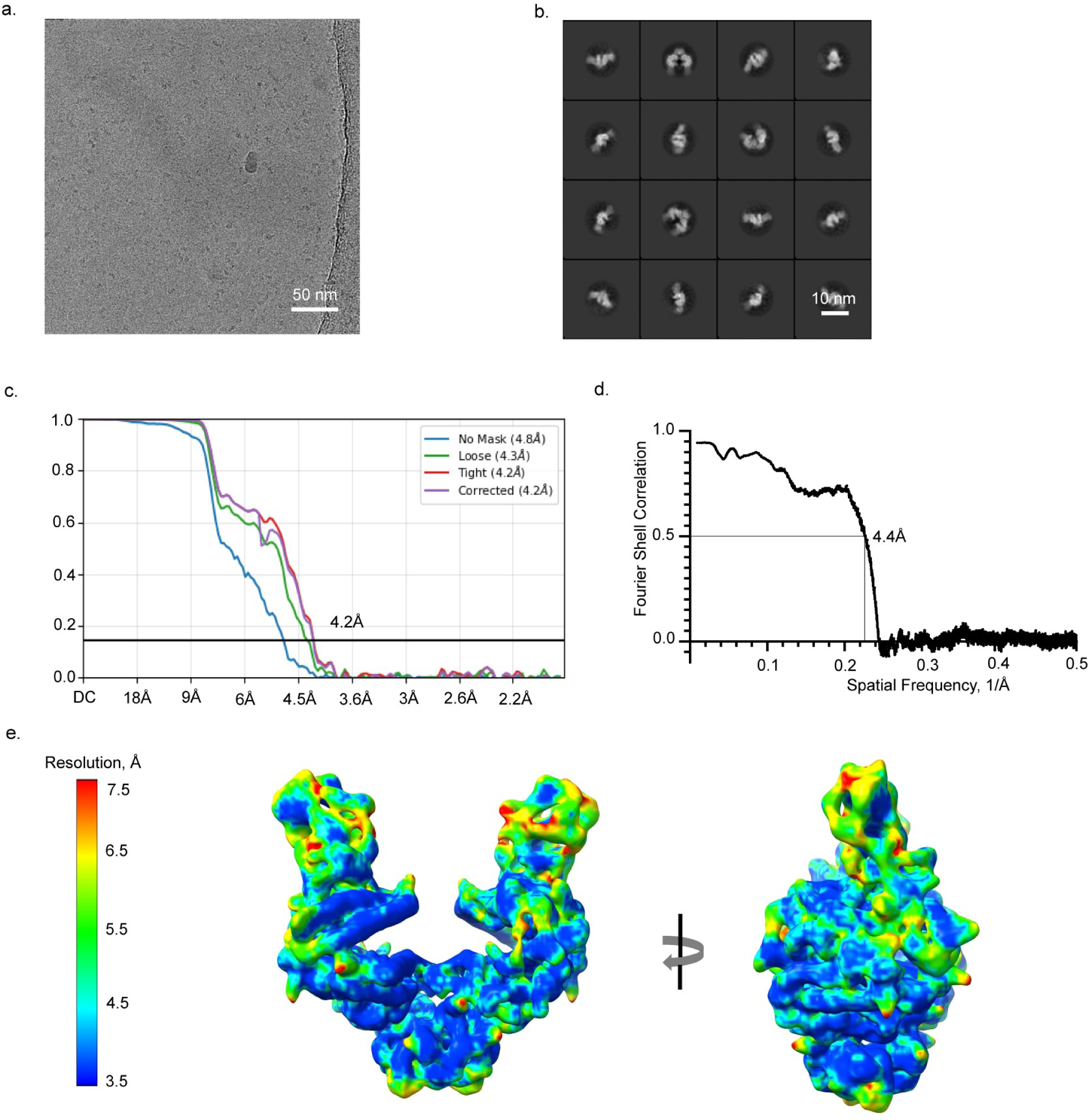
Validation of the cryo-EM map of *P. abramus* ACE2 in complex with HKU5-RBD. **a,** Representative micrograph. **b,** Representative high-resolution 2D class averages. **c,** Gold-standard resolution data generated by cryoSPARC. At the 0.143 threshold, the resolution is 4.2 Å. **d,** Fourier shell correlation curve between the map and the atomic model. **e,** Results of local resolution analysis using ResMap. The cryo-EM map is colored according to local resolution.

**Extended Data Fig. 6.**
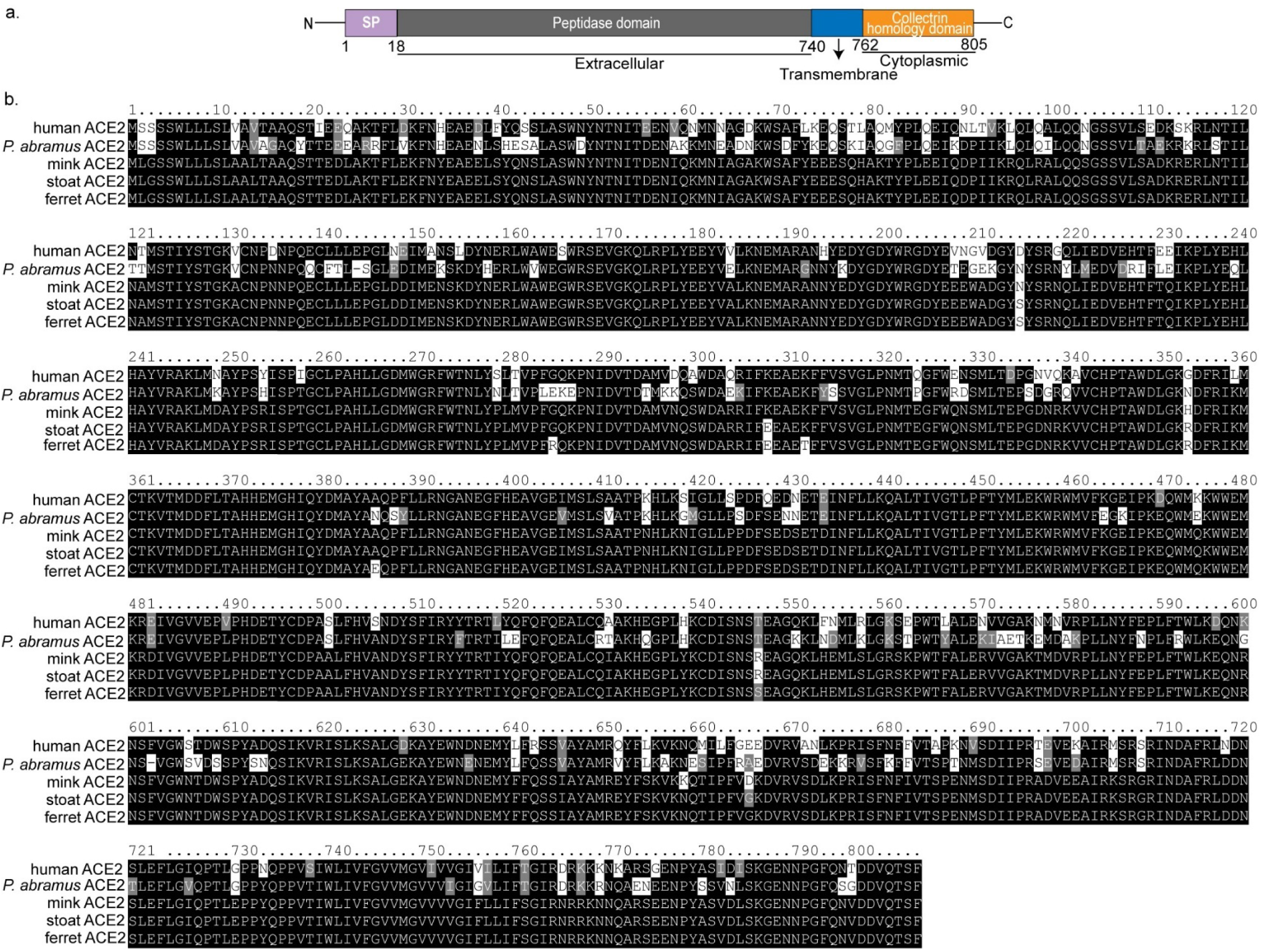
Schematic and alignment of featured ACE2 sequences. **a**, Schematic overview of ACE2 domains. **b,** Amino acid alignment of featured ACE2 sequences.

**Extended Data Fig. 7.**
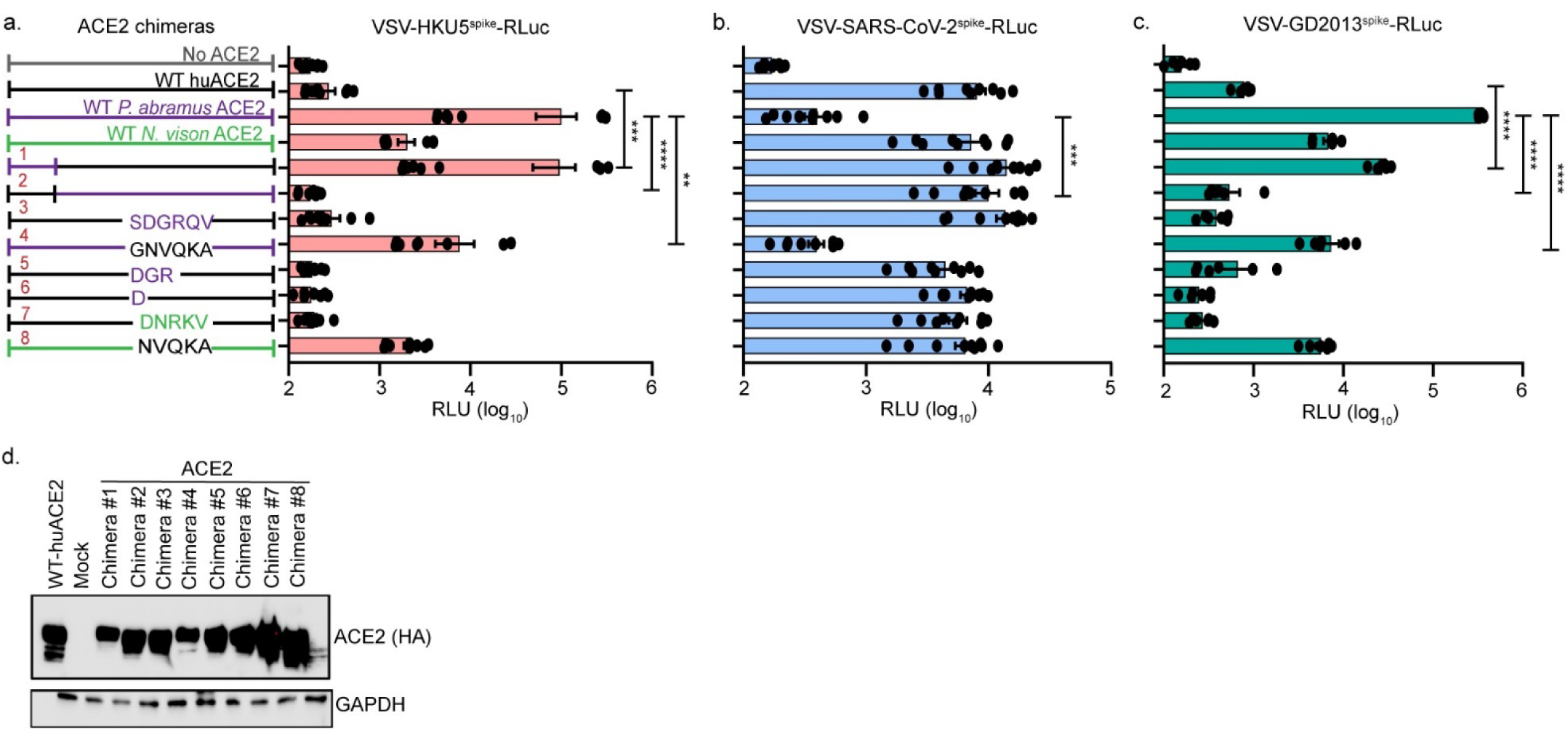
The N-terminal domain of *P. abramus* ACE2 mediates binding of HKU5 and GD2013 RBD. **a-c**, Schematic of the ACE2 chimeras between human, *P. abramus*, and *N. vison* ACE2 (left panel). The ACE2 chimeras were transiently transfected into HEK-293T cells and then infected with VSV-HKU5-^spike^RLuc (a), VSV-SARS-CoV-2^spike^RLuc (b), or VSV-GD2013^spike^-RLuc (c). **d,** HEK-293T cells were transfected with ACE2 chimeras containing a C-terminal HA tag and expression was confirmed by Western blot. Data are pooled from two independent experiments each done in triplicate. Statistical analysis was performed using two-tailed unpaired Student’s *t*-tests. The data presented are mean ± s.e.m (technical triplicate). *\*P<0.05,**P<0.01, ***P<0.001,****P<0.0001*.

**Extended Data Fig. 8.**
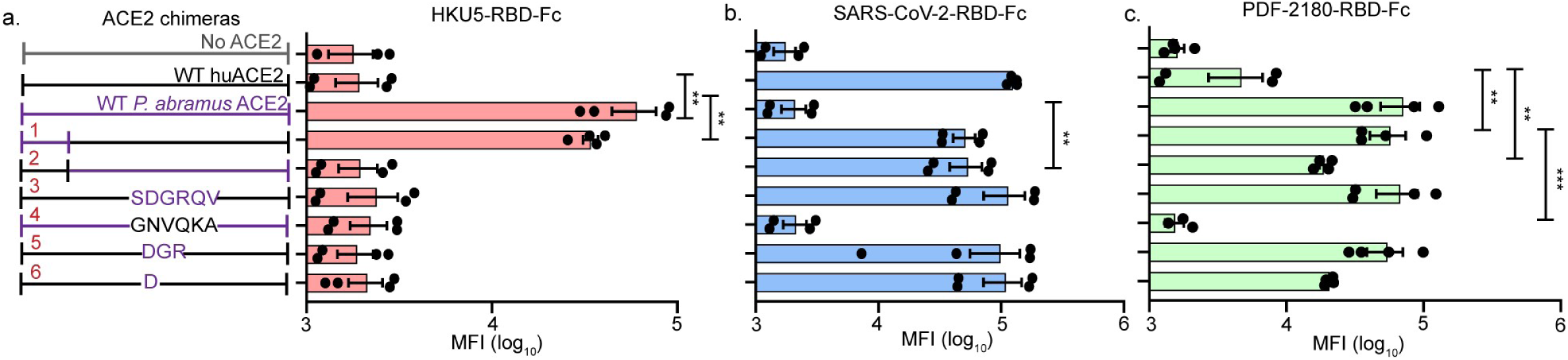
The N-terminal domain of *P. abramus* ACE2 is necessary and sufficient for HKU5 entry. **a-c**, Flow cytometry analysis of HKU5-RBD-Fc, SARS-CoV-2-RBD-Fc, and PDF-2180-RBD-FC binding to HEK-293T cells expressing the indicated WT (wild-type) and ACE2 chimeras. Mean fluorescence intensity (MFI) is shown. Data are pooled from two independent experiments each done in duplicate. Statistical analyses were performed using two-tailed unpaired Student’s *t*-tests and one-way ANOVA. Data are mean ± s.e.m. *\*P<0.05,**P<0.01, ***P<0.001,****P<0.0001*.

**Extended Data Fig. 9.**
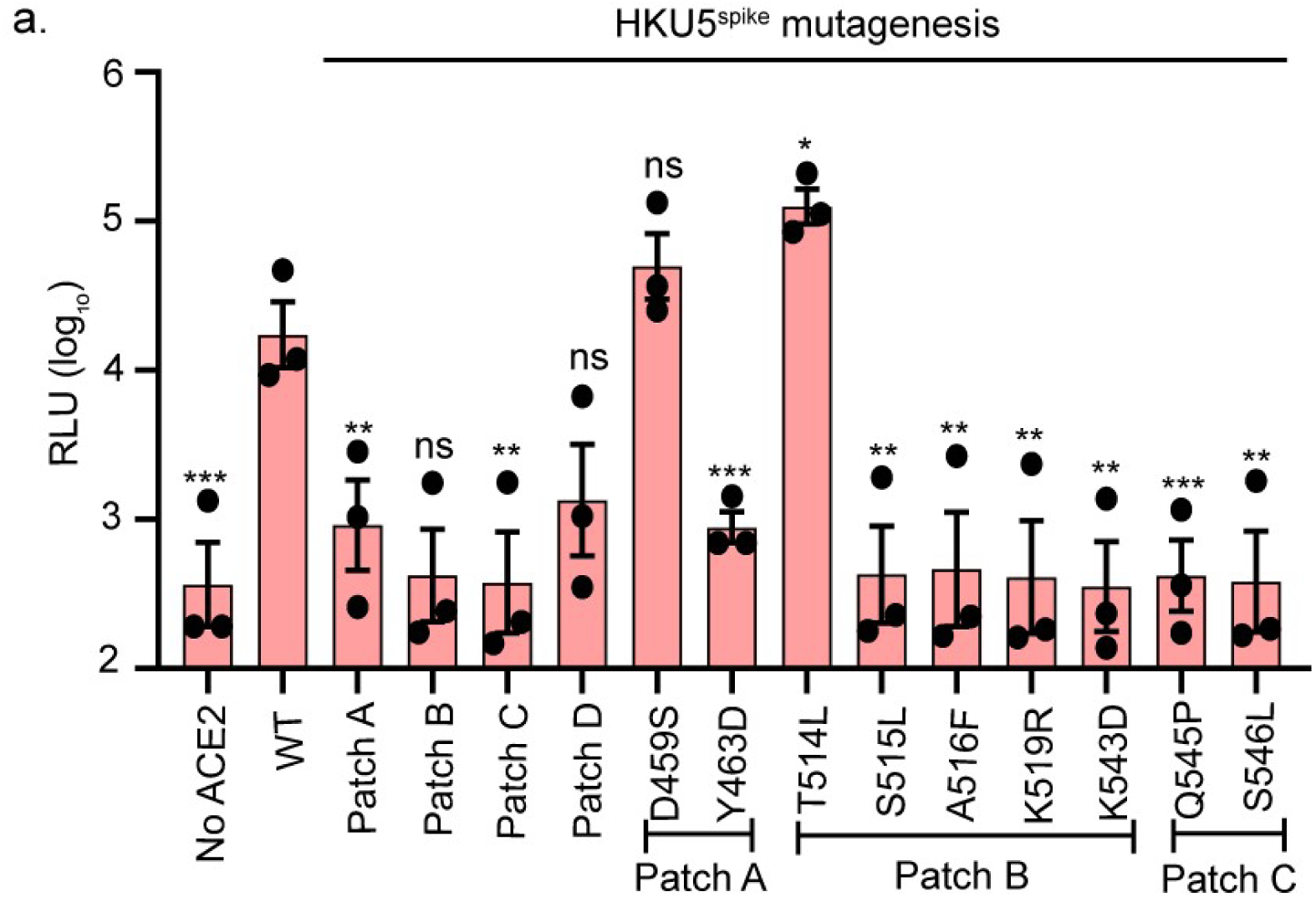
HKU5 spike mutations reveal determinants of *N. vison* ACE2-mediated entry. **a**, Four patches of substitutions on the HKU5-RBD were generated and tested for infectivity on HEK-293T cells transiently expressing *N. vison* ACE2. All four HKU5 patches of substitutions along with individual substitutions Y463D, S515L, A516F, K519R, K543D, Q545P, and S546L restricted *N. vison* ACE2 use. Data are mean from three independent experiments each done in triplicate. Statistical analyses were performed using two-tailed unpaired Student’s *t*-tests. Data are mean ± s.e.m. *\*P<0.05,**P<0.01, ***P<0.001*.

**Extended Data Fig. 10.**
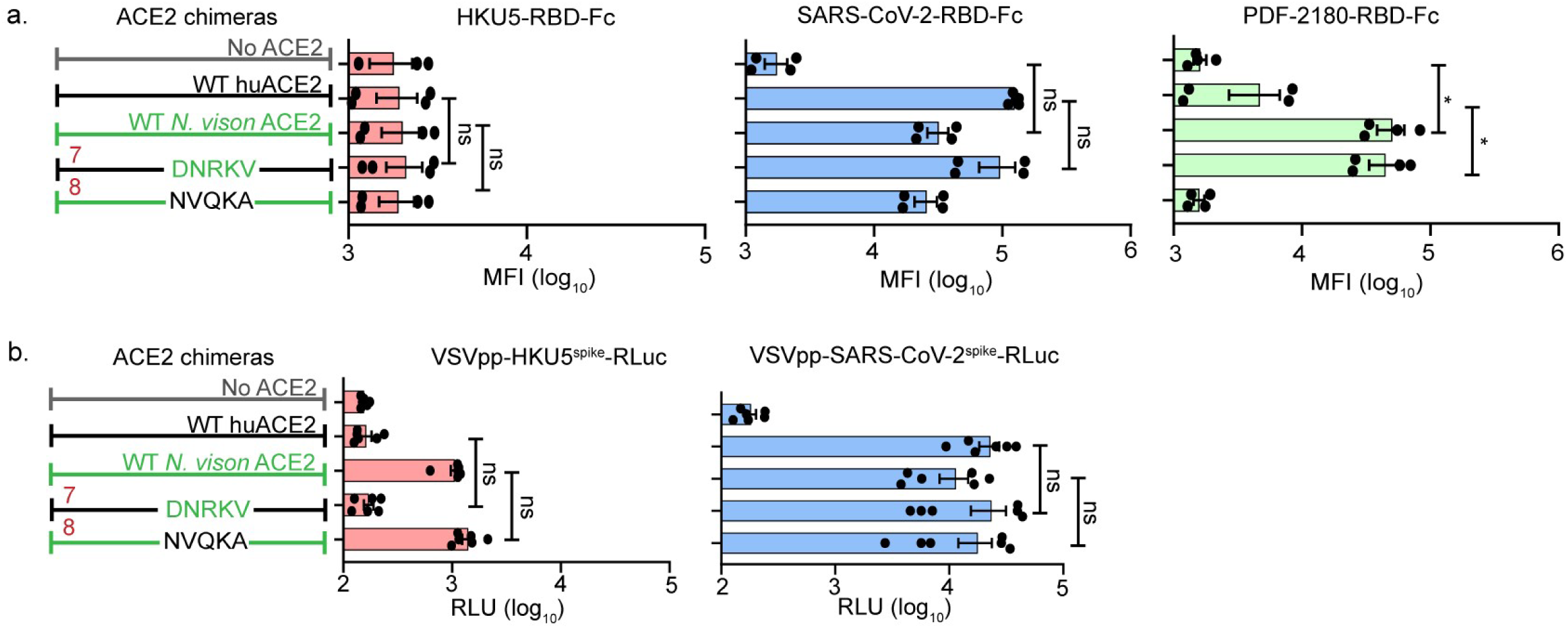
*N. vison* ACE2 residues 338-342 are neither necessary nor sufficient for HKU5 entry. **a**, Schematic of ACE2 chimeras between human and *N. vison* ACE2s (left). Flow cytometry analysis of HKU5-RBD-Fc and SARS2-RBD-Fc binding to HEK-293T cells expressing the indicated WT and ACE2 chimeras. The mean fluorescence intensity (MFI) was calculated. Residues 338-342 are necessary and sufficient for efficient binding of PDF-2180 in the context of *N. vison* ACE2 but not for HKU5 or SARS-CoV-2. **b,** ACE2 constructs were transiently transfected into BHK-21 cells, then infected with the VSV-SARS-CoV-^spike^RLuc or the VSV-HKU5-^spike^RLuc. Residues 338-342 were neither necessary nor sufficient for HKU5 entry in the context of *N. vison* ACE2. Data in (a) are pooled from two independent experiments each with two technical replicates. Data in (b) are mean from two independent experiments each done in triplicate. Statistical analyses were performed using two-tailed unpaired Student’s *t*-tests and one-way ANOVA. Data are mean ± s.e.m. *\*P<0.05*.

**Supplementary Table 1.**
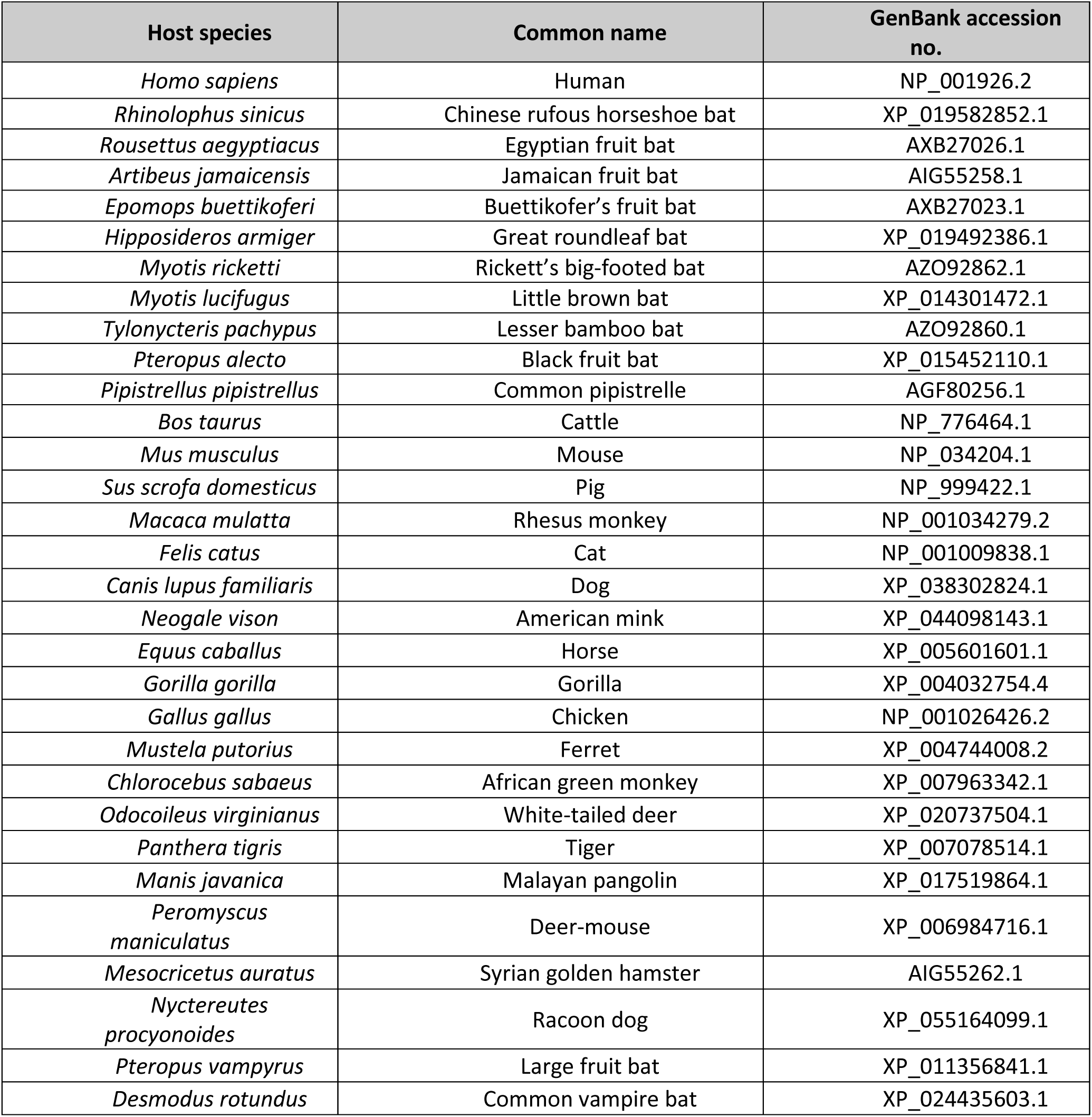
List of the 31 DPP4 orthologs used in this study.

**Supplementary Table 2.**
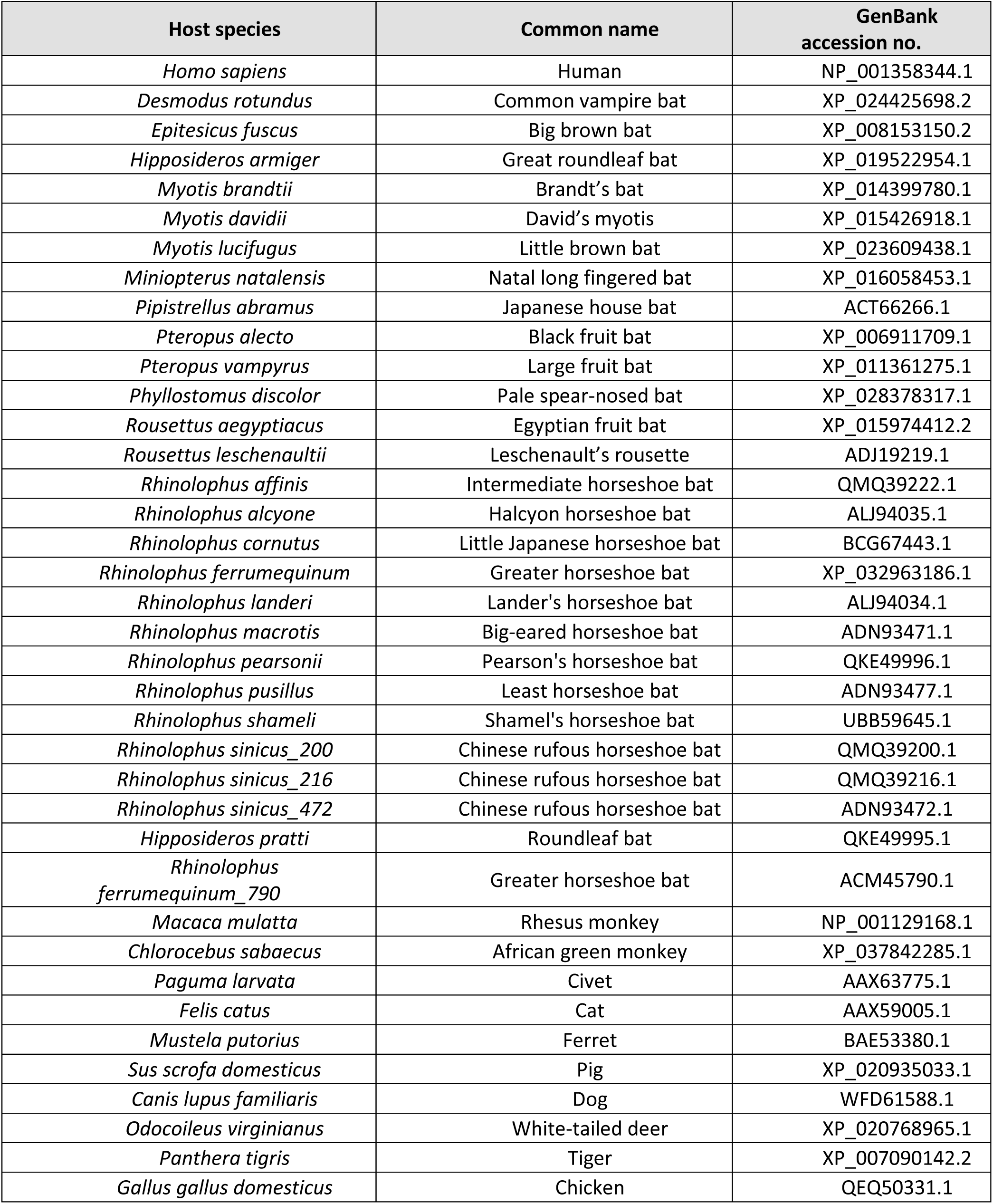
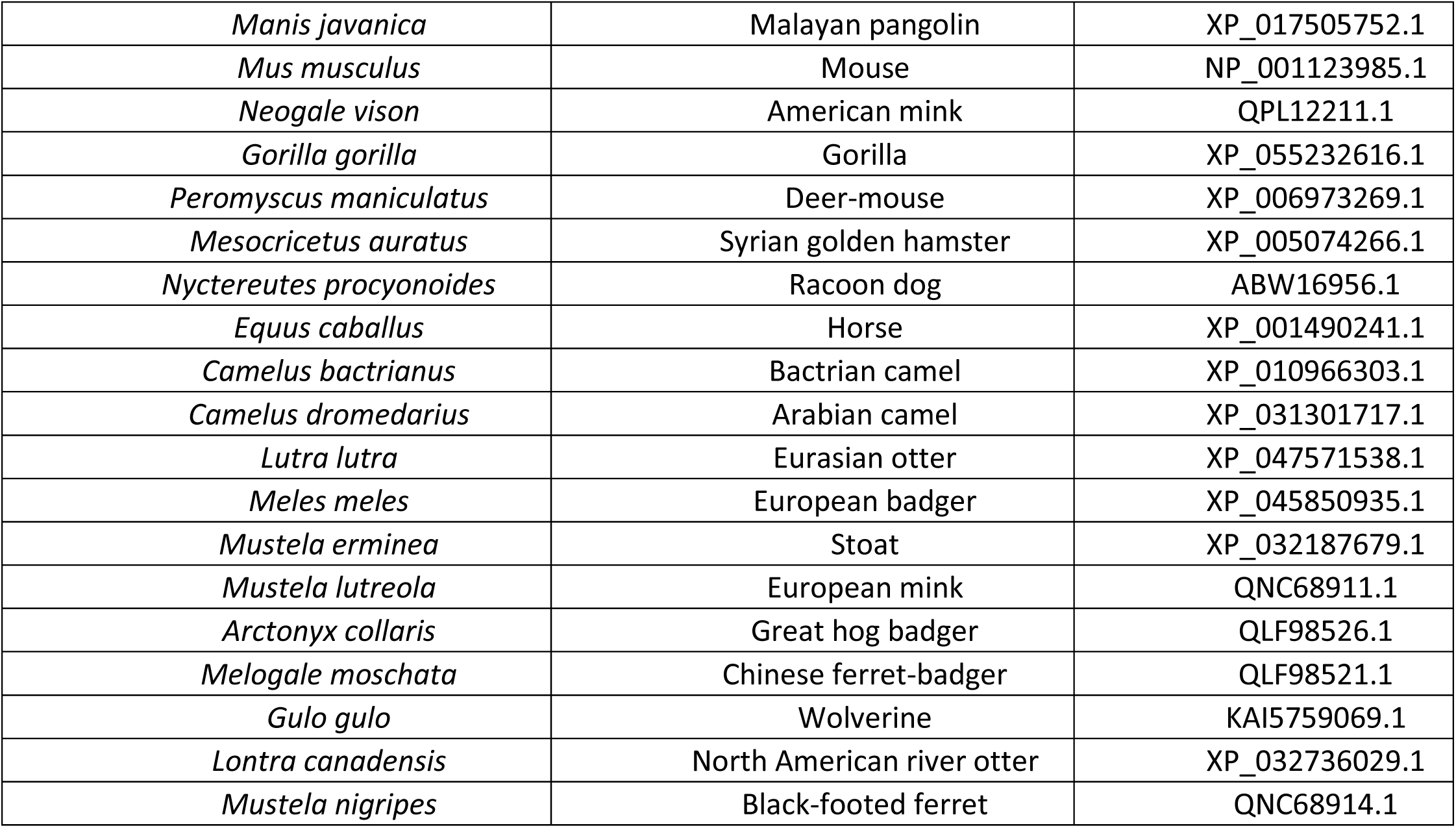
List of the ACE2 orthologs used in this study.

**Supplementary Table 3.**
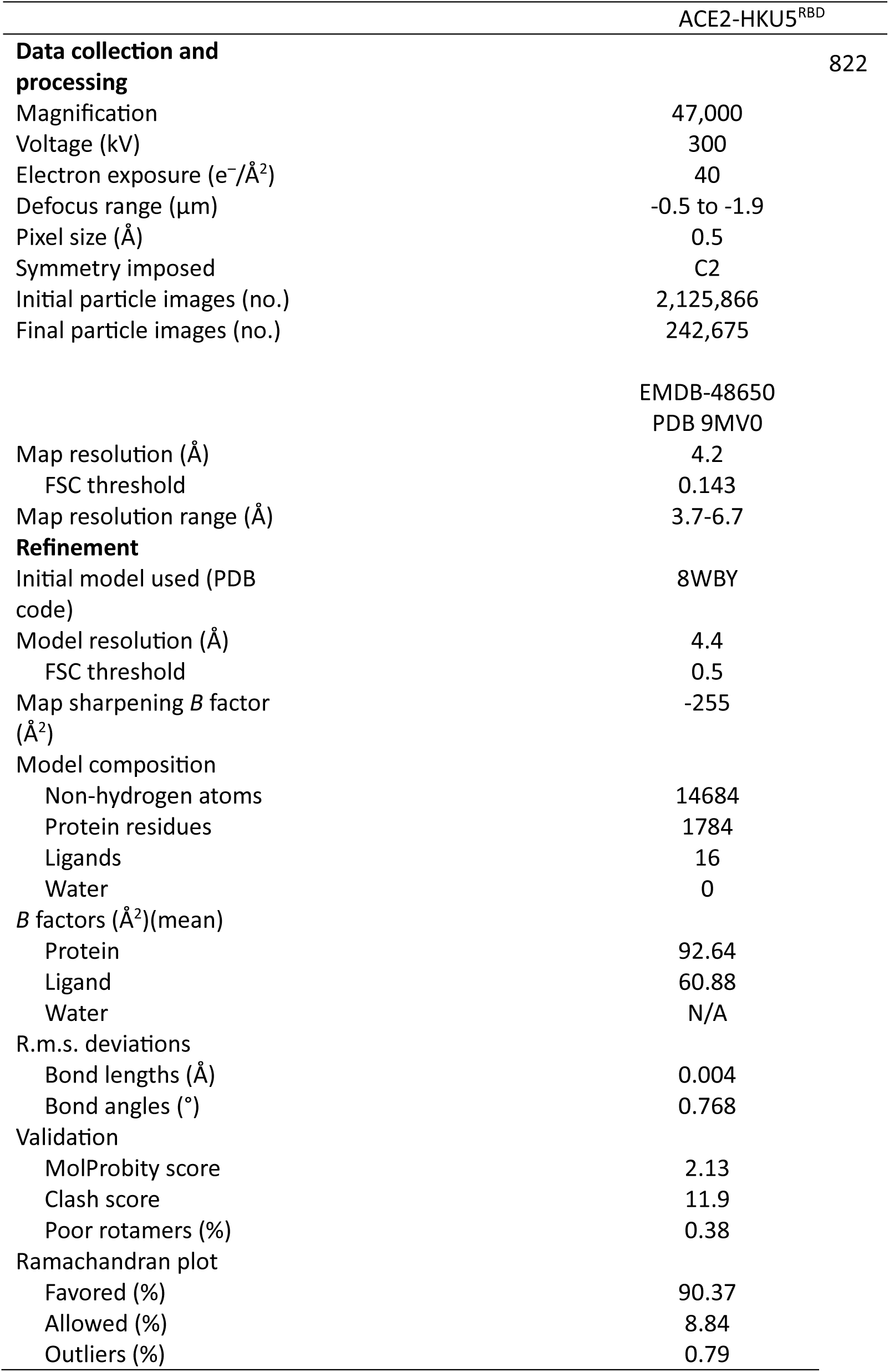
Cryo-EM refinement statistics.

**Supplementary Table 4.**
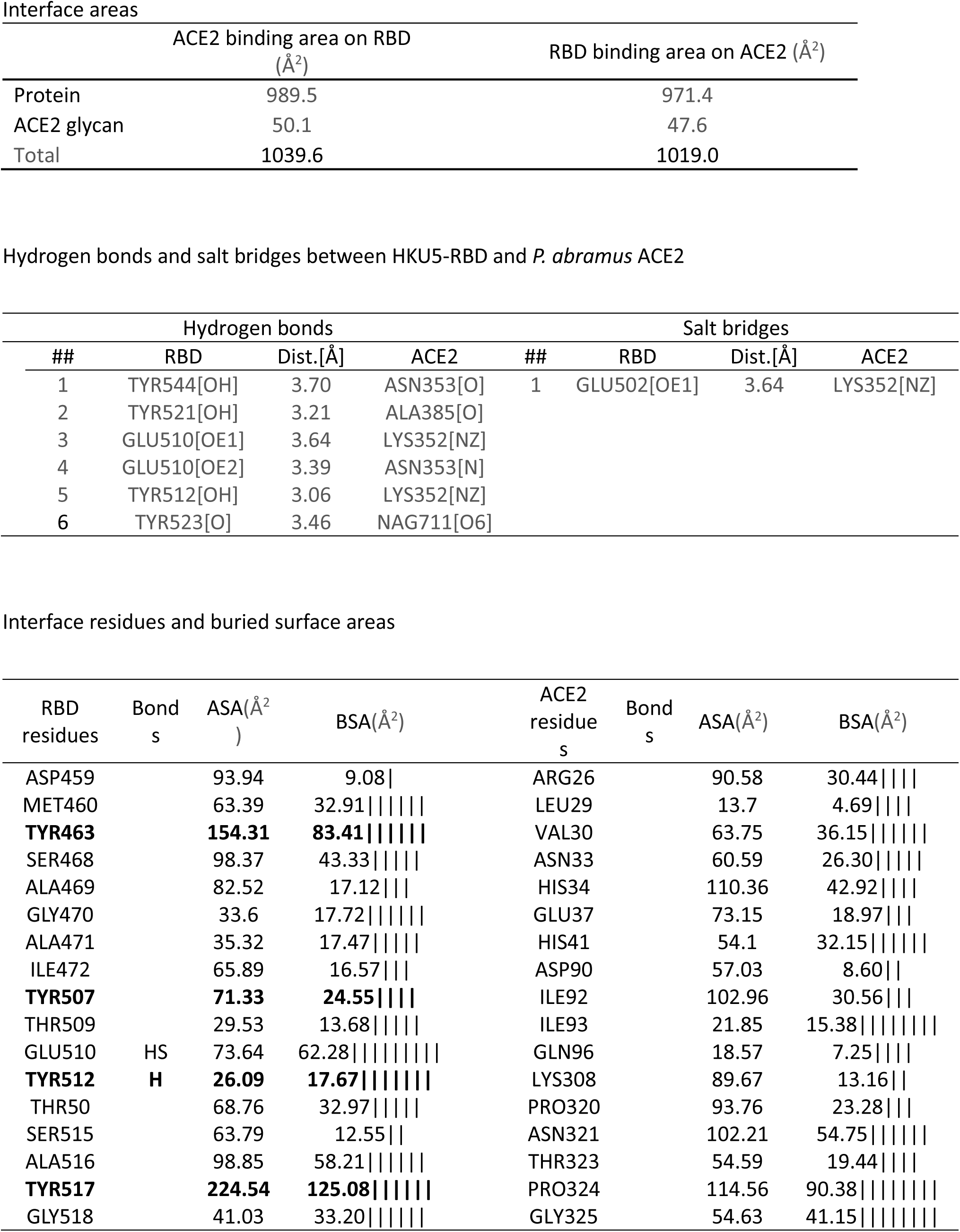
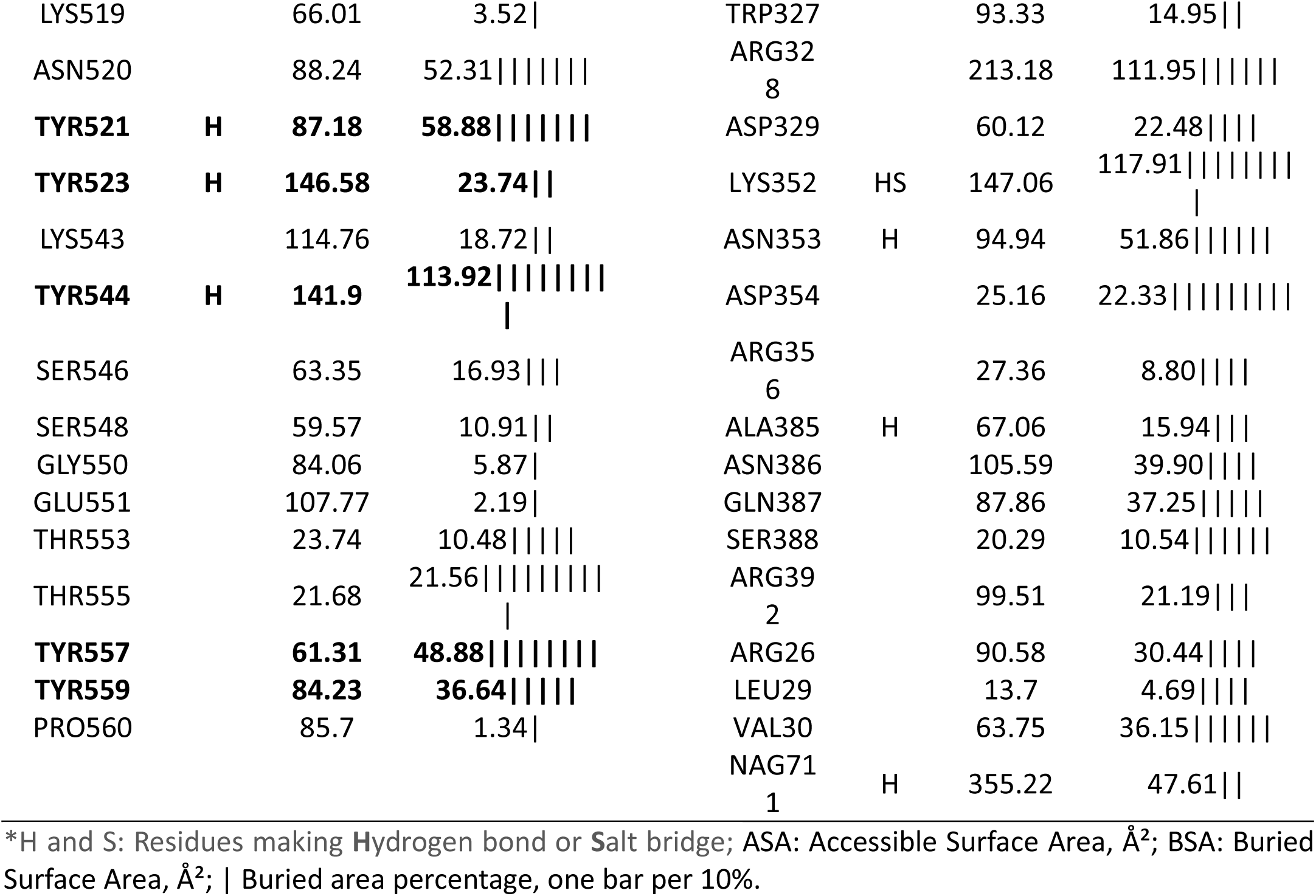
Interactions between HKU5-RBD and *P. abramus* ACE2.

